# A Re-Analysis of an Existing *Drosophila melanogaster* Dataset Reveals a New Set of Genes Involved in Post-Mating Response

**DOI:** 10.1101/2024.04.10.588867

**Authors:** Chloe J. Bennett, Rodolfo Aramayo

## Abstract

RNA sequencing (RNA-seq) is a commonly used method to identify changes in gene expression between two conditions. The analysis of RNA-seq output is complicated, with the possibility of getting different results from the same raw data. We developed and deployed four parallel pipelines to reanalyze an existing dataset of two female *Drosophila melanogaster* tissue types before and after mating. The *Drosophila* post-mating response (PMR) is a well-characterized suite of changes that occur after mating, accompanied by a flux in gene expression. In comparing our study with the previous analysis of this dataset, we find our results to be more stringent, though we do identify a number of significant genes not found before. We also found variation among our own separate experiments, with gene-to-transcript isoform number and index building playing important roles in outcome. Finally, we identified a set of genes found by our pipeline that were not identified by the previous study and proposed potential roles for these genes in post-mating biology. Together, this work presents a critique of current RNA-seq analysis techniques and proposes multiple workflow adjustments that can increase the sensitivity, specificity, and stringency of differential gene expression studies.

## INTRODUCTION

Differential Gene Expression (DGE) Analysis is a powerful tool when examining differences between two experimental, metabolic, and/or developmental conditions. Changes in gene expression underlie many behavioral and physiological phenotypes and being able to reliably identify which specific genes are being up- or downregulated can tell us more about the pathways that control these outcomes.

In the past, DGE analysis was solely accomplished using DNA microarray technologies (Bier et al., 2008). Briefly, microarrays rely on the hybridization of fluorescently labeled cDNA to known DNA sequences that have been printed on a chip, with relative fluorescent intensity indicating expression levels of the printed genes (Schena et al., 1995). Compared to expensive and time-consuming Sanger sequencing-based methods, or limited quantitative PCR methods, microarrays were a major breakthrough for studying genome-wide expression quickly and at a relatively low cost.

However, microarray DGE analysis is not without pitfalls. One major limitation of microarrays is the inability to uncover novel transcripts; only the predetermined genes printed on the chip can be detected. Additionally, due to hybridization kinetics and saturation, the fluorescent readouts from microarray chips can only provide an indirect, relative measurement and absolute expression levels cannot be determined (Held et al., 2006). DNA microarrays also generally lack the resolution to differentiate between genes with high sequence homology, though careful design of the chip can allow for detection of splice variants and alternative transcript isoforms (Castle et al., 2003). Microarrays are also plagued by a high false positive rate, requiring confirmation by follow-up experiments such as Northern blotting or PCR-based methods to ensure specificity (Wang et al., 2006).

With the advent of next-generation sequencing, RNA sequencing (RNA-seq) has eclipsed microarrays as the main modality for studying differential gene expression. RNA-seq first appeared in the published literature in 2008, with a dramatic increase in the number of publications referencing the technique, peaking in 2020 (7,272 manuscripts, as of this writing, according to PubMed). In contrast, publications referencing “*microarray*” and “*differential*” without the term “*RNA-seq*” have decreased since their 2012 peak, reaching a low of 1,242 in 2019. These trends in publications demonstrate the adoption of RNA-seq technology and the movement away from microarrays as a tool to study DGE.

A typical RNA-Seq experiment is generally composed of five distinct steps: experimental design, RNA isolation, library construction, sequencing, and differential gene expression analysis. When designing RNA-seq-based DGE experiments, a critical choice is deciding on sequence coverage and the number of replicates to use, with some groups suggesting a minimum of six biological replicates (Schurch et al., 2016). Total RNA is isolated from samples, enriched for the RNA type of interest (dependent on experiment), and converted to a cDNA library with ligated adapters (Wang et al., 2009). After sequencing comes the complicated task of analysis. Reads must undergo quality control measures before being aligned to the genome or transcriptome, quantified, and subjected to statistical tests to determine comparative levels of expression between samples (reviewed in Koch et al. (2018)).

RNA sequencing offers the potential to directly quantify all RNA present in a sample. Compared to array-based methods, RNA-seq is more able to detect transcripts with low levels of expression, identify biologically relevant isoforms, and detect splice variants and single nucleotide polymorphisms (Zhao et al., 2014). Another realized benefit of sequencing over arrays is the ability to reanalyze the resulting data later with different software and updated reference genomes/transcriptomes.

The output of an RNA-seq experiment are sets of genes that are either up- or downregulated in one condition compared to another condition (i.e. control). Inferences can be made based on the qualities of these resulting gene lists using downstream analytic tools. Single candidate genes can be further studied, but often a more general approach is used to identify classes of genes that are over- or underrepresented within the differentially expressed (DE) genes. One of the most widely used methods for categorizing DE genes is Gene Ontology (GO) analysis. Genes are assigned GO terms based on their associated molecular function, biological process, or cellular component (Consortium, 2008, 2019). Genes of interest within DGE results can be identified by determining which GO terms have significantly different representation within the set.

In this work, we selected to analyze a previously-generated dataset studying the post-mating response (PMR) in *Drosophila melanogaster* (Fowler et al., 2019). The PMR encompasses the stereotyped behavioral and physiological changes, and the accompanying dramatic fluctuations in gene expression, that occur in female flies after mating. Specifically, we aimed at testing if we could reproduce previously obtained results and, perhaps, identify new molecular targets associated with the PMR. While our main objective was to test the replicability and reproducibility of the previously published analysis, we also wanted to test if we could simplify and improve the computational pipeline. In their study, Fowler et al. (2019) sought to determine changes in expression of both coding and non-coding genes in two tissue types of male and female flies after mating, as well as to demonstrate the sex-based differences in PMR gene expression. The fully paired ncRNA and mRNA datasets from 16 samples allowed us to ask questions about gene regulation in the PMR, giving us the opportunity to explore different approaches for DGE analysis at both mRNA and sRNA level.

## METHODS

### Data Preparation

FASTQ files corresponding to messenger RNA (mRNA) and non-coding-RNA (ncRNA) were all subjected to a stringent quality control essentially as described (Blank et al., 2017; Maitra et al., 2020). Briefly, adapters were trimmed from the 51 nt raw reads. Duplicate reads were removed. Reads were scanned for bases with a FASTQ quality score of >20 and/or for the presence of ’N’s’, at which point they would be truncated and retained only if at least 15 nt. The reduction in read numbers after quality control for mRNA and sRNA are shown in Supplementary Tables S1 and S2, respectively (Aramayo, 2022).

### Mapping

#### Index Construction

The Indexes used in this work were prepared as follows (Fig. 1 and File S1, for details see Aramayo (2022): Using a FASTA file corresponding to the *Drosophila melanogaster* transcriptome (release 28), we generated the *Bowtie Indexed mRNAs* and the *Salmon Indexed mRNAs* indexes (Aramayo, 2022). The *Drosophila melanogaster* mRNA transcript sequences present in this transcriptome FASTA file were then clustered at 100% Identity and 100% Overlap with CD-HIT-EST (Li et al., 2001, 2002; Li and Godzik, 2006; Li et al., 2008; Huang et al., 2010; Niu et al., 2010; Fu et al., 2012; Li et al., 2012; Aramayo, 2022) to generate a *Drosophila melanogaster* clustered transcriptome FASTA file that was then used to generate the *Bowtie Indexed Clustered mRNAs* and the *Salmon Indexed Clustered mRNAs* indexes (Fig. 1A)(Aramayo, 2022). For the Canonical analysis, we used the transcript IDs corresponding to the Canonical mRNA transcripts to extract their corresponding FASTA sequences from the *Drosophila melanogaster* transcriptome (release 28), FASTA file, to generate a *Drosophila melanogaster* Canonical transcript sequences only FASTA file. This file was used to generate both the *Bowtie Indexed mRNAs* and the *Salmon Indexed mRNAs* indexes (Aramayo, 2022).

**Figure 1.**
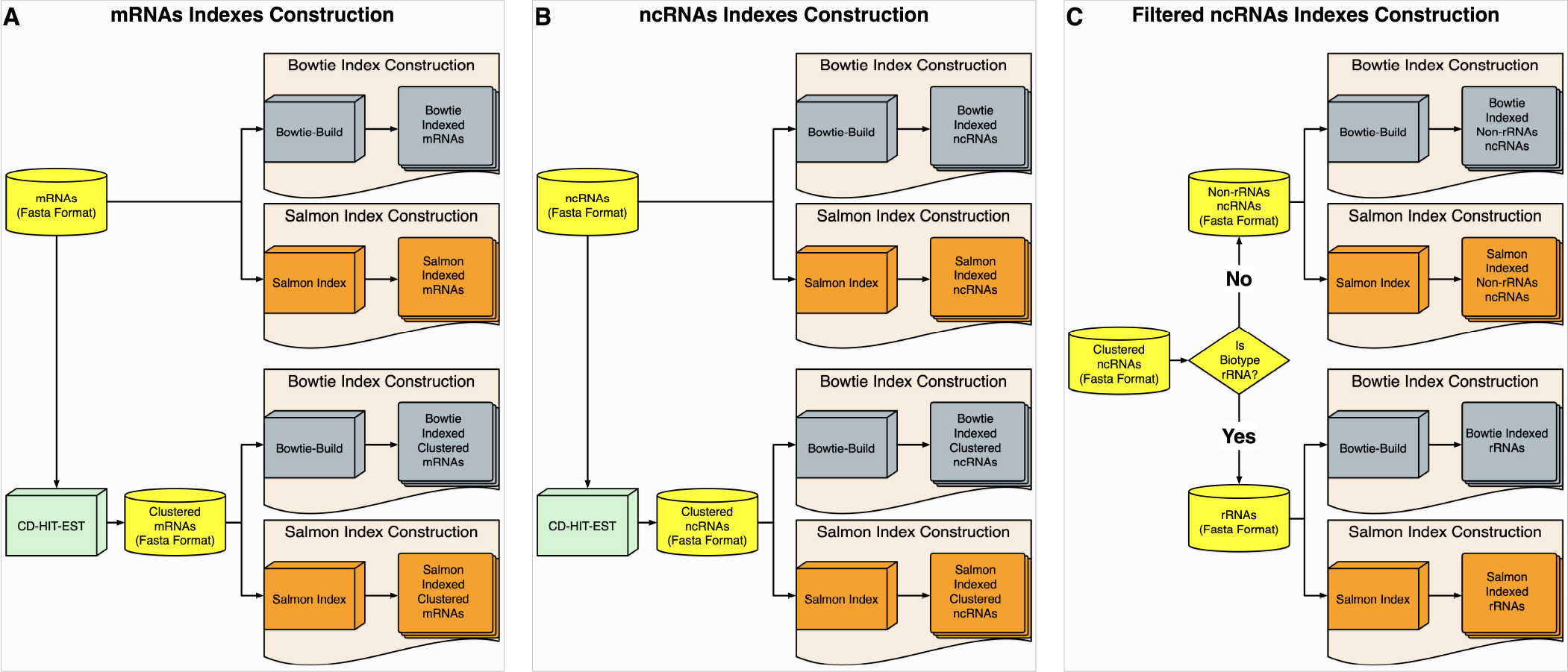
**Index creation pipelines**. (A) mRNA indexes were built using either Bowtie or Salmon either directly or after clustering similar sequences with CD-HIT-EST. (B) ncRNA index construction before or after clustering. (C) Clustered ncRNAs were subjected to a filtering step using indexes of rRNA alone and non-rRNA ncRNA alone.

Similarly, using a FASTA file corresponding to the *Drosophila melanogaster* ncRNA (release 28), we generated the *Bowtie Indexed ncRNAs* and the *Salmon Indexed ncRNAs* indexes. The *Drosophila melanogaster* ncRNA transcript sequences present in the ncRNA FASTA file were then clustered at 100% Identity and 100% Overlap with CD-HIT-EST (Li et al., 2001, 2002; Li and Godzik, 2006; Li et al., 2008; Huang et al., 2010; Niu et al., 2010; Fu et al., 2012; Li et al., 2012) to generate a *Drosophila melanogaster* clustered ncRNA transcriptome FASTA file that was then used to generate the *Bowtie Indexed Clustered ncRNAs* and the *Salmon Indexed Clustered ncRNAs* indexes (Fig. 1B)(Aramayo, 2022).

Finally, the transcript IDs corresponding to the clustered *Drosophila melanogaster* ncRNA transcripts were separated into ribosomal RNA (rRNA) and non-rRNA. Transcript IDs corresponding to non-rRNA ncRNA were then used to extract their corresponding FASTA sequences to generate a file containing their sequences. Transcript IDs corresponding to rRNA were similarly used to extract their corresponding FASTA sequences to generate a file containing the rRNA transcript sequences. The FASTA file corresponding to the non-rRNA was then used to generate both the *Bowtie Indexed Non-rRNAs ncRNAs* and the *Salmon Indexed Non-rRNAs ncRNAs* indexes (Fig. 1C). Similarly, the FASTA file corresponding to the rRNA was then used to generate both the *Bowtie Indexed rRNAs ncRNAs* and the *Salmon Indexed rRNAs ncRNAs* indexes (Fig. 1C). *Bowtie* indexes were constructed using default command parameters. *Salmon* indexes were constructed with the *–kmerLen 9* flag on.

#### Bowtie Mapping

Both mRNA and ncRNA reads were mapped with Bowtie version 1.2.3 (Langmead et al., 2009), against indexes described above. Bowtie mapping was performed with the *-a -v 3 –best and –strata* flags on.

#### HISAT2 Mapping

When needed, mRNA and ncRNA reads were also mapped against the *Drosophila melanogater* genome using the *Drosophila melanogaster* genome (release BDGP6.28), with HISAT2 version 2.1.0 (Kim et al., 2019, 2015; Pertea et al., 2016; Zhang et al., 2021b)), using the following parameters: *–dta –dta-cufflinks –new-summary –min-intronlen 40 –max-intronlen 200000*.

### Salmon Pseudo Mapping and Quantification

Mapped reads were quantified using Salmon version 1.3.0 and 1.9.0 (Patro et al., 2017). Salmon was run with the following parameters: *quant –seqBias –gcBias –libType U –numBootstraps 100*. We used a *Drosophila melanogaster GTF* annotation file version (BDGP6.28.47) (see File S1).

### Differential Expression Analysis

When needed, the package wasabi was used to process Salmon output for downstream analysis. Sleuth was used to identify transcripts with significantly different abundances between virgin and mated female abdomens (VFAB x MFAB) and virgin and mated female head-thorax (VFHT x MFHT) (Pimentel et al., 2017).

### Differential Expression Analysis

After the initial alignment/pseudo-counting, the resulting FASTQ files were prepared for Sleuth (Pimentel et al., 2017) analysis using wasabi, and then processed with Sleuth (Pimentel et al., 2017), using custom made one-liners (see File S1, for details) (Aramayo, 2022). Significantly regulated transcripts as declared by the Wald test in Sleuth (Pimentel et al., 2017), were then identified and processed. Tables generated by Sleuth containing the lists of statistically significant transcripts were calculated by the Wald Test at a False Discovery Rate of 0.05. To translate the transcript IDs in these tables to gene IDs, we compiled two relational tables: zzMasterTableChrGeneTransName_01 and zzMasterTableChrGeneTransName_02. Both of these tables associate gene names with gene IDs, transcript IDs, and transcript biotypes (File S2) (Aramayo, 2022). These tables were constructed by extracting the sequence annotation present in the FASTA headers of the genomes files: *Drosophila_melanogaster.BDGP6.28.cdna.all.fa* (mRNA) and *Drosophila_melanogaster.BDGP6.28.ncrna.fa* (ncRNA), and processing this information using the commands summarized in File S2. The resulting tables were then used to classify the Sleuth data tables and to translate Gene IDs to Transcript IDs and transcript biotypes. The information present in the Sleuth tables was then processed using the commands summarized in the File S2 file and from this we extracted the Gene, Transcript, and Biotype IDs present in the different tables. All commands issued are described in File S2. We calculated the overlap of the DE genes from our analysis to those of Fowler et al. (2019) using the commands described in the sections called “Calculating Overlaps” of File S2 (Aramayo, 2022).

### IGV Visualization

When needed, SAM files were converted/sorted into BAM files and displayed with IGV (Busan and Weeks, 2021; Robinson et al., 2017; Busan and Weeks, 2017; Coletta et al., 2012; Thorvaldsdottir et al., 2013; Robinson et al., 2011). We identified a set of constitutively expressed genes to serve as visual controls (Corrales et al., 2017; Lü et al., 2018). These genes are: *Vps35 (*FBgn0034708), *eIF3i/Trip1 (*FBgn0015834), *ATPsynB (*FBgn0019644), *Cyp1 (*FBgn0004432), *GstD1 (*FBgn0001149), *Rap2l (*FBgn0283666), *CG13220 (*FBgn0033608), *Robl (*FBgn0024196), *RpL32 (*FBgn0002626), *RpL13 (*FBgn0011272), *RpS20 (*FBgn0019936), and *eIF2 (FBgn0261609)*. To facilitate visualization, only genes less than or equal to 4000 bp in length were considered.

## RESULTS AND DISCUSSION

### Experimental Rationale: Background and Strategy

In their study, Fowler et al. (2019) performed RNA-seq on total RNA isolated from groups of 50 male or female, virgin or mated, abdomens (Ab) or head/thoraxes (HT). They used a blocking oligo for their ncRNA samples to reduce the adaptor ligation, and thus detection via sequencing, of 2S rRNA. They obtained a total of two biological replicates for each sample type, resulting in 32 total samples (2 replicates x 2 sexes x 2 tissue types x 2 mating conditions x 2 RNA types). After determining DE mRNA and ncRNA between various samples, they performed miRNA target prediction using the TargetScan algorithm (Lewis et al., 2005; Riffo-Campos et al., 2016). They also performed Gene ontology enrichment analysis using GOrilla (Eden et al., 2009). Although this study is undoubtedly important in that it offered the first direct comparison of post-mating gene expression in both sexes of *Drosophila*, as well as being the first one to examine post-mating miRNA expression between sexes of any species, it is not without issues. The study’s resulting mRNA and ncRNA data were both analyzed using divergent computational pipelines. Additionally, only miRNAs were identified within the ncRNA data. Perhaps the largest issue with the study was the choice to use only two replicates per condition, limiting the statistical power and ability to draw strong conclusions. While being powerless to correct for the lack of replicates, we selected this study due to its other attributes and aimed at testing the possibility of developing a single computational pipeline to re-quantify both mRNA and ncRNA datasets simultaneously. We hypothesized that a reanalysis of the data could, while confirming previously obtained results, reveal the role of new genes/pathways in the PMR. In summary, our analysis was aimed at testing the reproducibility/replicability of the study while exploring the feasibility of analyzing mRNA and sRNA data in a single computational pipeline.

DGE analysis can evaluate statistically significant differential regulation at either gene- or transcript-level. Such evaluation normally uses a single computational pipeline. Fowler et al. (2019) performed their analysis at the gene level. To test if we could identify a new set of DE genes, not detected by Fowler et al. (2019), we designed four different computational pipelines (or experiments) aimed not only at detecting transcript level (in addition to gene level) differential expression, but also at comparing the performance and differences between various alignment and filtering techniques while evaluating those results with a single pseudoalignment and quantification routine (see Figs. 2 and 3). To accomplish this, we restricted our re-analysis to comparing DGE observed only between virgin and mated female abdomens (VFAB x MFAB) and virgin and mated female combined heads and thoraxes (VFHT x MFHT) conditions.

**Figure 2.**
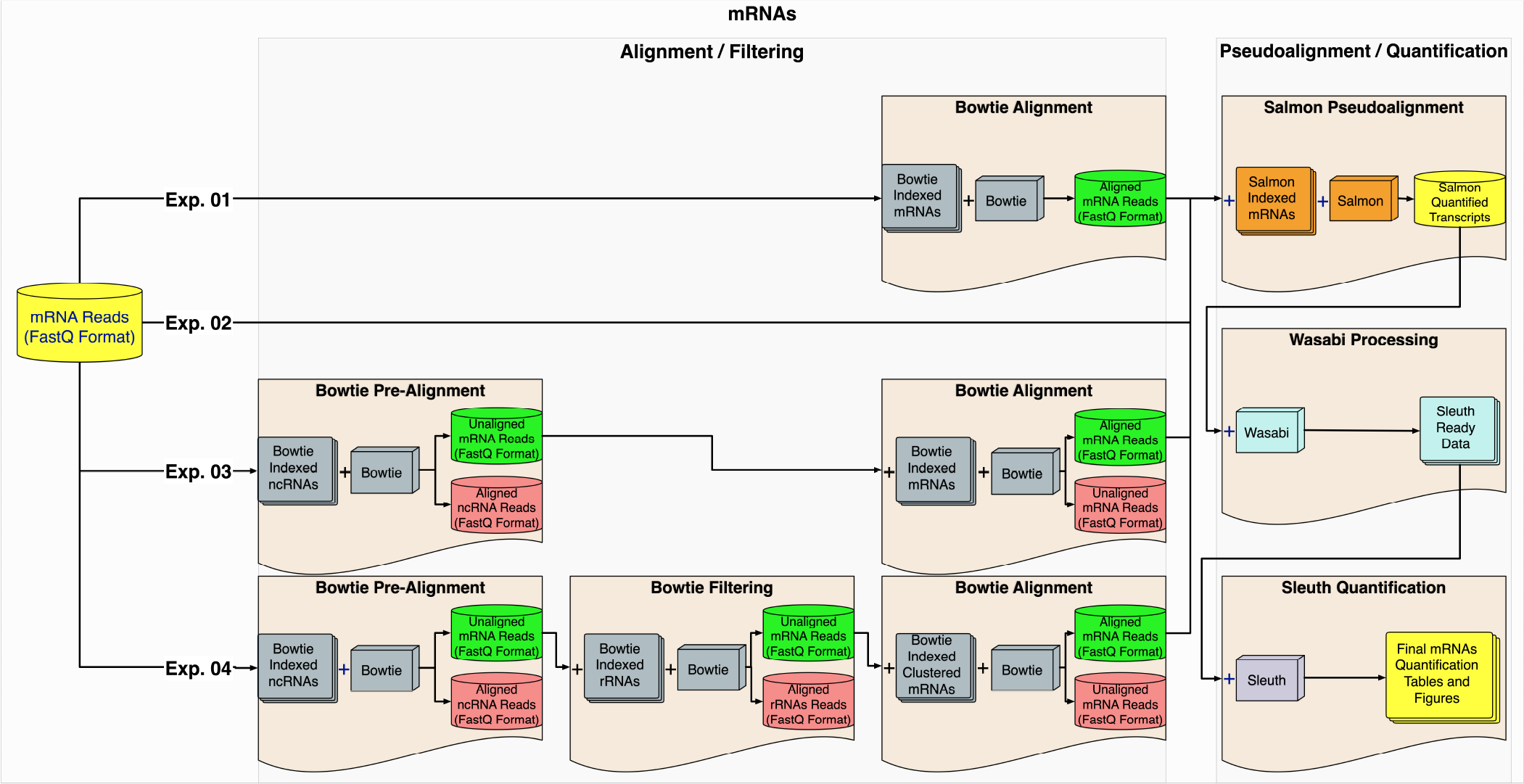
**mRNA processing pipelines**. (Exp. 1) mRNA reads were aligned with Bowtie and fed into Salmon/Wasabi/Sleuth processing pipeline. (Exp. 2) mRNA reads were fed directly into Salmon/Wasabi/Sleuth processing pipeline. (Exp. 3) mRNA reads were subjected to a pre-filtering step by aligning them to the ncRNA index, before processing them as in Exp. 1. (Exp. 4) mRNA reads were subjected to a further filtering step by pre-aligning them to both the ncRNA and rRNA indexes, before being processed as in Exp. 1.

**Figure 3.**
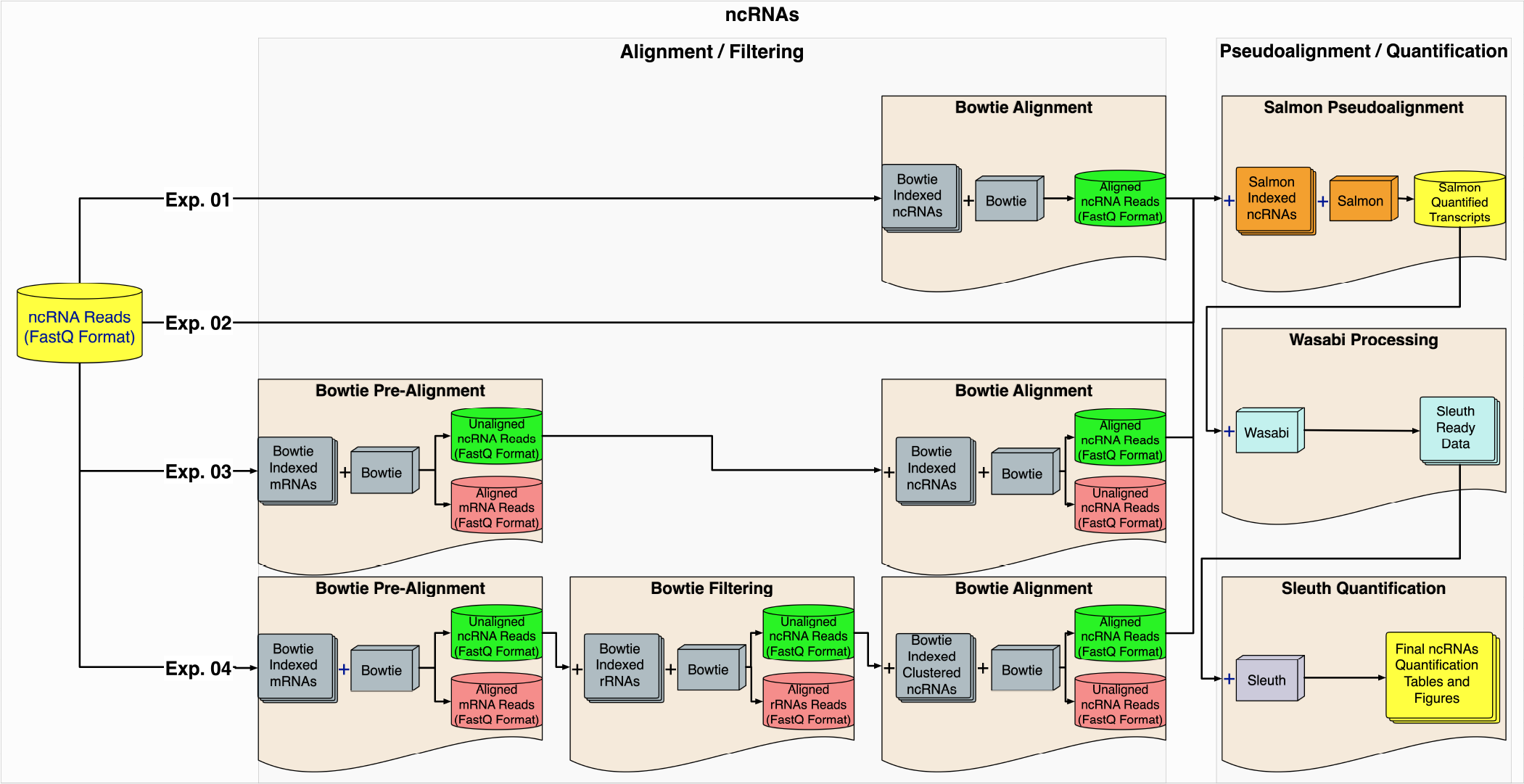
**ncRNA processing pipelines**. (Exp. 1) As with the mRNAs, the ncRNA reads were processed using four pipelines. (Exp. 1) ncRNA reads were aligned with Bowtie and fed into Salmon/Wasabi/Sleuth processing pipeline. (Exp. 2) ncRNA reads were fed directly into Salmon/Wasabi/Sleuth processing pipeline. (Exp. 3) ncRNA reads were subjected to a pre-filtering step by aligning them to the mRNA index, before processing themas in Exp. 1. (Exp. 4) ncRNA reads were subjected to a further filtering step by pre-aligning them to both the mRNA and rRNA indexes, before being processed as in Exp. 1.

In all cases, the quality-controlled mRNA or ncRNA reads corresponding to either the VFAB x MFAB or VFHT x MFHT conditions were processed by four different pipelines (or experiments). In Experiment 1, the mRNA or ncRNA reads were first aligned with Bowtie using either *Bowtie Indexed mRNAs* index (for mRNAs, Fig. 2), or *Bowtie Indexed ncRNAs* index (for ncRNAs, Fig. 3). The resulting FASTQ files containing either the Bowtie-aligned mRNA or ncRNA reads were then sent to quantification using the Salmon/Wasabi/Sleuth pipeline. In Experiment 2, the mRNA or ncRNA reads were directly evaluated by the Salmon/Wasabi/Sleuth pipeline (see Figs. 2 and 3). In Experiment 3, the mRNA or ncRNA reads were pre-aligned with Bowtie using the *Bowtie Indexed ncRNAs* for mRNA reads or the *Bowtie Indexed mRNAs* for ncRNA reads. The objective of this pre-alignment was to remove potential contaminating ncRNA present in the mRNA reads, or mRNA present in the ncRNA reads. The resulting unaligned reads were then aligned and quantified exactly as described for Experiment 1. Finally, for Experiment 4, in addition to the pre-alignment performed and described in Experiment 3, the resulting pre-aligned reads were subjected to another filtering step aimed at removing potential rRNA reads present in either the mRNA or ncRNA data. Similar to Experiment 3, the resulting pre-aligned and rRNA-filtered reads were then processed as described for Experiment 1.

### Dataset Preparation, Pre-Alignment, and Filtering

Exps. 1 and 2 were aimed at determining if pre-mapping the reads with Bowtie before their pseudoalignment/quantification changed the results obtained by directly feeding those QC-processed FASTQ files into the Salmon/Wasabi/Sleuth pipeline. As such, other than performing stringent Quality Control (see Materials and Methods), FASTQ files were directly quantified as described in Fig. 2 (mRNA) or Fig. 3 (ncRNA). In contrast, for Exps. 3 and 4, FASTQ files were subjected to an additional alignment. With this pre-alignment, Exp. 3 was aimed at detecting potential contamination of ncRNA reads in mRNA datasets or mRNA reads in ncRNA reads. Similarly, Exp. 4 aimed at not only detecting potential contamination of ncRNA in mRNA reads or mRNA in ncRNA reads, but also at testing and quantifying the presence of contaminating rRNA reads in all datasets (see Exps. 3 and 4 in Figs. 2 and 3).

Pre-aligning mRNA reads to a ncRNA Index identified *∼*2% of the mRNA reads as being ncRNA reads. The vast majority of the mRNA reads (i.e., *∼*98%), behaved as expected (Fig. 4). Approximately 89.65% of those reads aligned to the annotated mRNAs, and only *∼*10% of the reads failed to align to either ncRNA or mRNA indexes (see Exp. 3, Fig. 4). We failed to detect the contamination of ncRNA corresponding to rRNA in the mRNA reads (see Exp. 4, Fig. 4).

**Figure 4.**
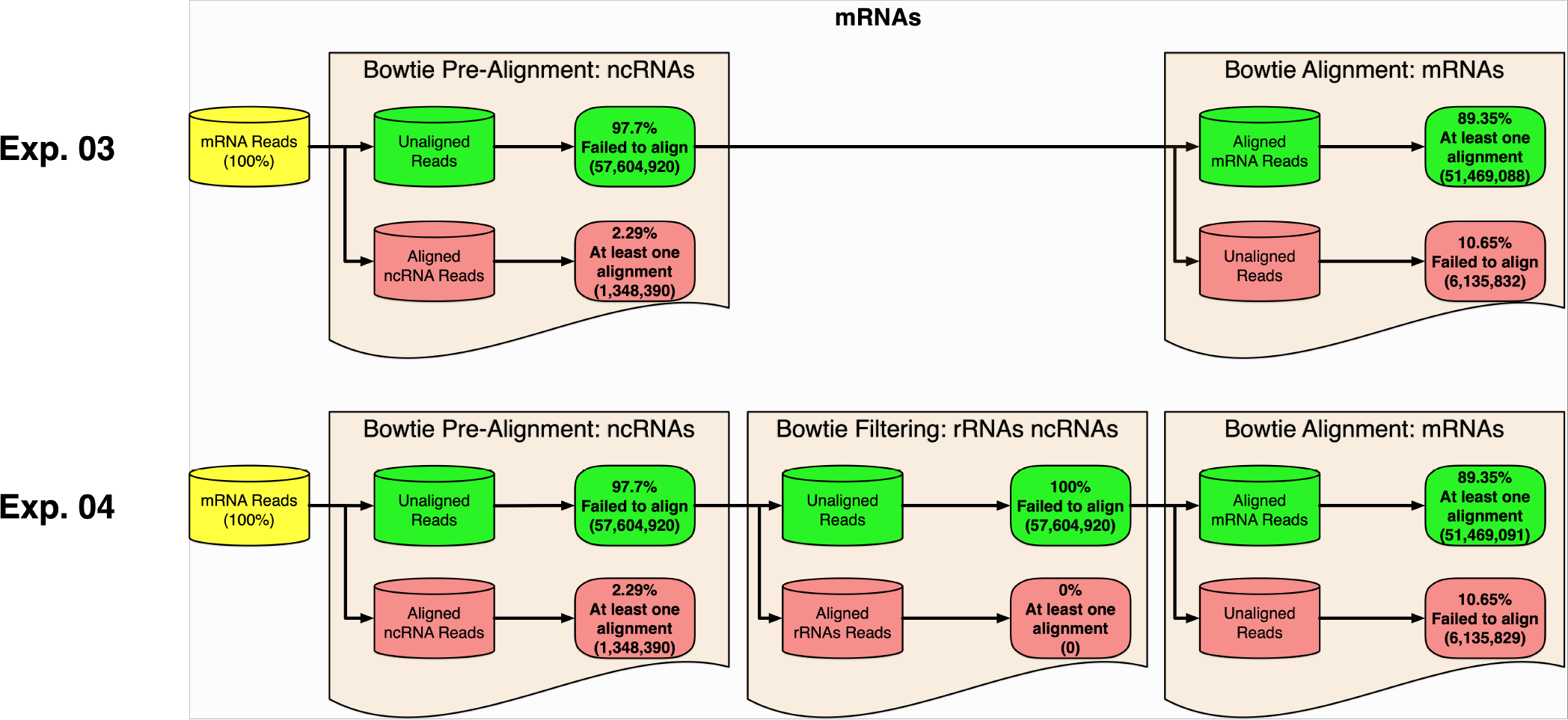
**Pre-alignment and filtering of mRNA reads**. For Exp. 3 and Exp. 4, pre-alignment of the mRNA reads to the ncRNA index revealed 2% of the mRNA reads were ncRNA contamination. No rRNA-specific contamination was detected. As expected, the majority of the reads that failed to align to the ncRNA index (i.e., 97.7%), aligned to the transcriptome (i.e., 89%).

In contrast, pre-aligning ncRNA reads to a mRNA Index identified that *∼*64% of the ncRNA reads are in fact mRNA reads. Only *∼*36% of the ncRNA reads failed to align to the mRNA Index. Surprisingly, only *∼*16% of those *∼*36% reads aligned to an index corresponding to annotated *Drosophila* ncRNAs. Here, the vast majority of the reads (i.e., *∼*84%) failed to align to either mRNA or ncRNA indexes (see Exp. 3, Fig. 5). These experiments detected a *∼*2% contamination of rRNA within the ncRNA reads (see Exp. 4, Fig. 5).

**Figure 5.**
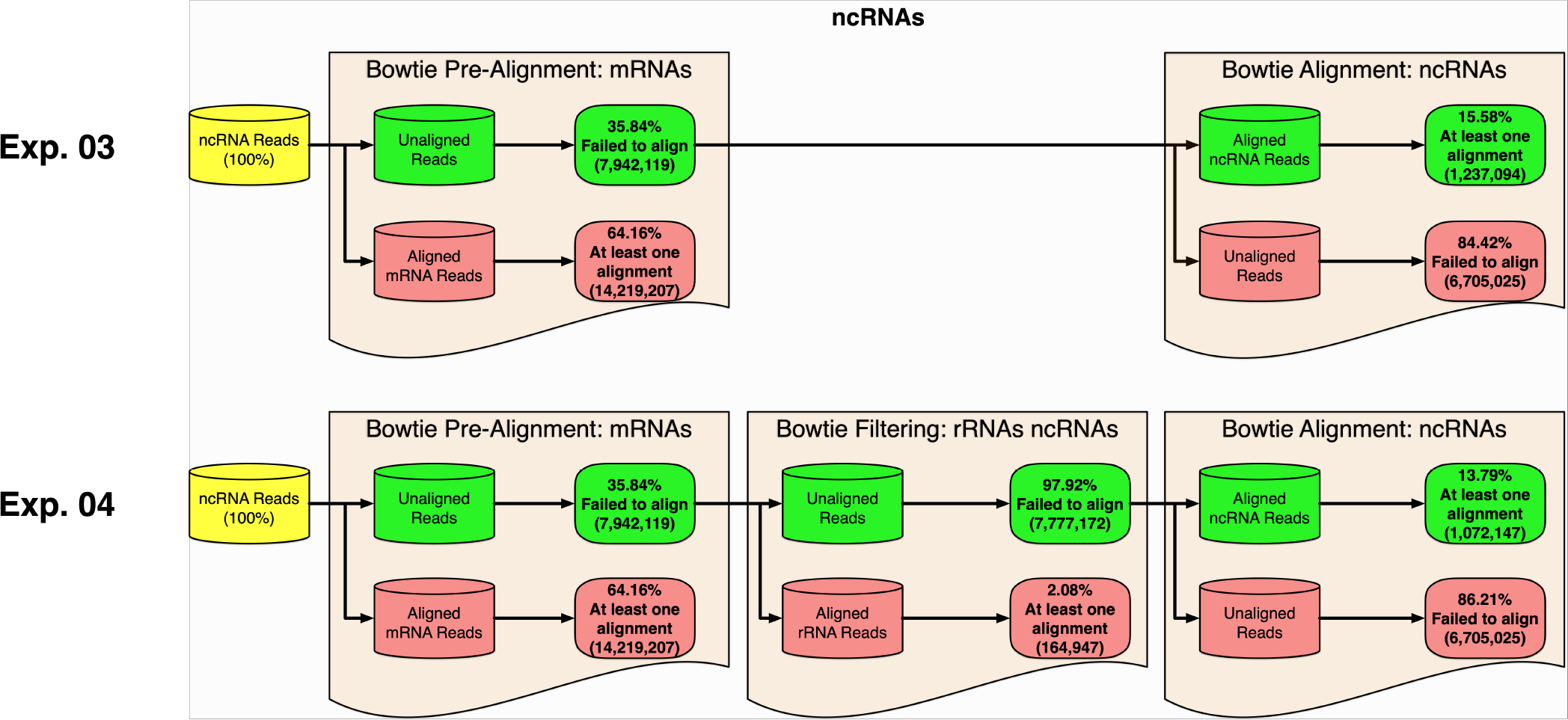
**Pre-alignment and filtering of ncRNA reads**. For Exp. 3 and Exp. 4, pre-alignment of the ncRNA reads to the mRNA index revealed 64% of the ncRNA reads were mRNA contamination. A 2% of the ncRNA reads were ncRNAs related to rRNAs. Only 15.58% and 13.79% of the reads were considered true ncRNA reads for Exp. 3 and Exp. 4, respectively.

These observations underscore the importance of determining the true composition of the data used in any DGE analysis, especially those involving ncRNA reads. We propose that a filtering strategy, as described in this work, be performed before any DGE analysis (see Discussion below).

### Differential mRNA Expression Analysis

#### Differentially Expressed mRNA (DE-mRNA) Analyses Produced An Overall Similar, But Not-Identical, Number of Significantly Regulated Transcripts

To determine the presence of significantly regulated mRNA and/or ncRNA transcripts identified in *all* of our experiments, we used the transcript IDs declared by the Sleuth’s Wald test to calculate the total number of *unique* hits and the number of *unique* hits present in one or more experiments (Table 1 and Figs. 6 and S1). As each gene ID is associated with one or more transcript IDs (based on the number of transcript variants), we calculated both the number of *unique* genes *and* transcripts detected by our experiments (See Tables 1 and 2).

**Figure 6.**
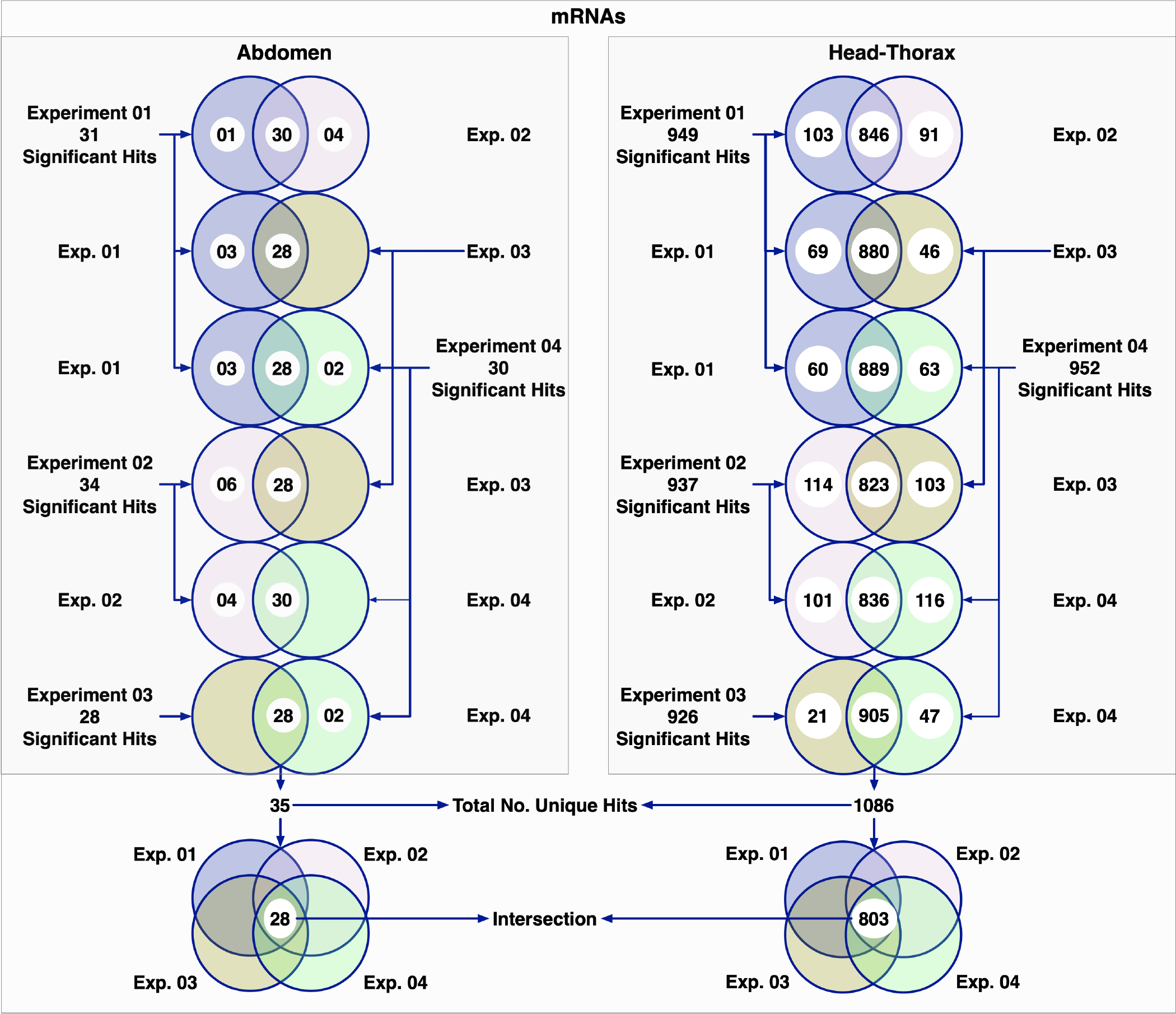
**Significant hit overlap between and among mRNA experiments**. The series of Venn Diagrams represent the number of significant genes detected when comparing two experiments (i.e., say Experiment A with Experiment B). The diagrams represent genes only present in Experiment A, only present in Experiment B or present in both Experiments A and B. The results of comparing of all four experiments to each other are displayed. Below, we summarize the results of the four experiments, which gave in Ab a total of 35 significantly DE transcripts, with 28 of those being identified by each of the four experiments. In HT, a total of 1086 significantly DE transcripts were observed, with 803 of those represented by each of the four experiments.

**Table 1.** Females: Virgin vs Mated. This table shows the number of mating-responsive transcripts found by each experiment for mRNA and ncRNA in both abdomen and head-thorax. The final two rows represent the number of transcripts identified across all four experiments (i.e. in any of the four) and the number of transcripts that were found in each of the four experiments (i.e. in all of the four).

| Experiment Name | Significant Transcripts<sup>1</sup> |  |  |  |
| --- | --- | --- | --- | --- |
|  | mRNAs |  | ncRNAs |  |
|  | Abdomen | Head-Thorax | Abdomen | Head-Thorax |
| Experiment 01: | 31 | 949 | 3 | 0 |
| Experiment 02: | 34 | 937 | 3 | 1 |
| Experiment 03: | 28 | 926 | 3 | 2 |
| Experiment 04: | 30 | 952 | 2 | 1 |
| **Total No. Unique Hits:** | **35** | **1086** | **5** | **3** |
| **Total No. Unique Hits in All 4 Experiments<sup>2</sup>:** | 28 (80%) | 803 (73.94%) | 1 (20%) | 0 |
| <sup>1</sup> Significant mRNA or sRNAs were calculated using the Wald Test at a False Discovery Rate of 0.05.<br><sup>2</sup> These numbers were calculated using the Total No. Unique Hits data. |  |  |  |  |

**Table 2.** Comparison of gene expression in All and Canonical experiments. In both Ab and HT, the Canonical experiment was able to detect significantly more DE genes than the All experiment. Of the 93 total gene IDs found in both experiments, only two were found by the All experiment alone, 66 were found just in the Canonical experiment, and 25 were identified by both experiments. The differences between experiments in HT showed similar proportions, but with overall higher numbers of DE genes. 2,278 unique gene IDs were identified across both experiments, 119 of which were from All, 1,525 from Canonical, and 634 from the overlap of both experiments.

|  | Significant Genes<sup>1</sup> |  |  |  |  |  |
| --- | --- | --- | --- | --- | --- | --- |
|  | All Transcripts | Canonical Transcripts | No. Combined Unique | Overlap (%) | Only All Transcripts | Only Canonical Transcripts |
| No. of Hits: | Abdomen |  |  |  |  |  |
|  | 27 | 91 | 93 | 25 (26.9%) | 2 (2.1%) | 66 (70.9%) |
| No. of Hits: | Head-Thorax |  |  |  |  |  |
|  | 753 | 2159 | 2278 | 634 (27.8%) | 119 (5.2%) | 1525 (66.9%) |
| <sup>1</sup> Significant mRNA were calculated using the Wald Test at a False Discovery Rate of 0.05. |  |  |  |  |  |  |

The number of differentially expressed transcripts detected by the different experiments described above was, in general, equivalent (see Fig. 6), but not identical. When performing a one-on-one comparison between the different experiments, we observed a high degree of overlap (Table 1 and Fig. 6). We also observed transcripts that were detected as significant in an experiment-specific manner. The identification of these experiment-specific hits raises the possibility that many of them might very well be false positives. Although combining the results of different experimental approaches has the potential of increasing the detection of true positives by reducing experimental noise (as we did in this work), there is demonstrable need for the development of computational pipelines that consider the existence of these experiment-specific hits.

Considering only hits that are *common* to all experiments (i.e. those detected by *all* experiments), we observed a total of 28 and 803 DE-mRNAs for Abdomen (Ab) and Head-Thorax (HT), respectively (see Table 1 and Fig. 6). The *∼*28 times higher number of hits observed in HT versus Ab condition, we believe, is directly related to intra- and inter-condition variation, as visualized in the principal component analyis (PCA) (Fig. S1 A and B).

We do not think these observations reflect a biological phenomenon; it seems that the closer two experimental replicates are to each other (as declared in both the PCA and dendogram behavior), the higher the number of differentially regulated mRNA transcripts that will be detected. The pair of replicates for the virgin condition in the HT dataset are closely related to each other (i.e., they behave as ideally expected), while the mated samples are less similar. The four samples represented in the Ab condition show much more variation, with a wide spread between replicates of the same condition and when comparing virgin to mated samples. We expect that less intra-condition and greater inter-condition variation would result in greater statistical power, raising the number of significant hits for samples meeting this described PCA behavior.

Although we observed a different number of DE-mRNAs for each experiment, overall the number of *common* DE-mRNAs detected was high (i.e., 80% for Abdomen and 74% for Head-Thorax, see Table 1 and Fig. 6). In contrast, we observed only 14 and 508 DE-mRNAs that were only seen only by either one of the four pipelines for Abdomen and Head-Thorax, respectively. If we consider these DE-mRNAs as false positives, then these experiments generated a total of 8.5% (3 out of 35) and 9.7% (106 out of 1086) false positives for Abdomen and Head-Thorax, respectively. For the rest of this work, we consider as significantly differentially regulated only those transcripts that were detected by all four experiments performed (i.e., 28 for Abdomen and 803 for Head-Thorax).

#### Differentially Expressed Gene (DE-Gene) Analyses Produced Different Numbers of Significantly Regulated Transcripts

We wanted to test the effect that using only one main canonical transcript per gene would have on the overall number of differentially regulated genes detected. The determination of DE-mRNAs takes into consideration the presence of *all* transcript isoforms corresponding to a given gene. As different isoforms generally have a high degree of sequence overlap, we reasoned that this could potentially interfere with any individual sequence read address assignment strategy used, thus introducing an unknown biased parameter in this equation. We hypothesized that by using only one main transcript per gene as defined by Ensembl (i.e. the canonical transcript), we would *normalize* our differential gene expression calculation, reducing any potential isoform-induced bias in the process. An Ensembl Canonical transcript is a single transcript chosen to represent a gene, defined using a series of different algorithmic parameters that take into consideration such factors as conservation, expression, and coding sequence length. We denote canonical transcripts in this text using a superscript C (i.e. FBtrX…X*^C^*). Files containing canonical transcripts were constructed as described in Materials and Methods and used to replace the equivalent files used in Exp. 4. We used the Exp. 4 pipeline strategy for this analysis because it was, in our view, the most stringent in terms of read quality control before the Psueudoalignment/Quantification step.

Results from this one-gene, one-canonical-transcript experiment gave very different results compared to previous experiments. This experiment identified 91 DE genes, as opposed to only 27 genes identified before, for Ab (see Tables 2 and 3). Similarly, for HT, this experiment identified 2,159 DE genes, compared to the 753 previously identified, for HT. This represents an increase of *∼*3.3 and *∼*2.9 times more genes detected for Ab and HT, respectively.

**Table 3.** Comparing gene expression across the current and 2019 studies. Using canonical transcripts to identify DE genes increased the similarity between our results and those of the 2019 study. The proportion of DE genes found only by the 2019 study decreased by at least half in both Ab and HT in the Canonical experiment compared to the All experiment. Both of our experiments were more stringent than the previous study in that they identified an overall lower number of significant hits.

|  | Significant Genes<sup>1</sup> |  |  |  |  |  |
| --- | --- | --- | --- | --- | --- | --- |
|  | This Study | 2019 Study | No. Combined Unique | Overlap (%) | Only This Study | Only 2019 Study |
| Abdomen |  |  |  |  |  |  |
| All Transcripts: | 27 | 130 | 134 | 23 (17.2%) | 4 (3%) | 107 (79.8%) |
| Canonical Transcripts: | 91 | 130 | 143 | 78 (54.5%) | 13 (9.1%) | 52 (36.3%) |
| Head-Thorax |  |  |  |  |  |  |
| All Transcripts: | 753 | 2048 | 2202 | 599 (27.2%) | 154 (7%) | 1449 (65.8%) |
| Canonical Transcripts: | 2159 | 2048 | 2668 | 1539 (57.7%) | 620 (23.2%) | 509 (19.1%) |
| <sup>1</sup> Significant mRNA were calculated using the Wald Test at a False Discovery Rate of 0.05. |  |  |  |  |  |  |

We compared the distribution of transcript IDs identified in this experiment to those detected in our previous, all-isoforms-included (All) experiments. We observed increased significant hits in the canonical experiment (see Fig. 7). Calculating the degree of overlapping hits detected between the Canonical experiment and the set of 28 unique significant hits detected in the All experiment, we found that *∼*71% (i.e. 20 out of 28) of the transcripts detected were also detected by the Canonical experiment. In contrast, in the reciprocal comparison, only *∼*22% (i.e. 20 out of 91) were detected by the Canonical experiment (Fig. 7). Overall, the degree of overlap detected was 26.9% for Ab and 27.8% for HT (see Table 2).

**Figure 7.**
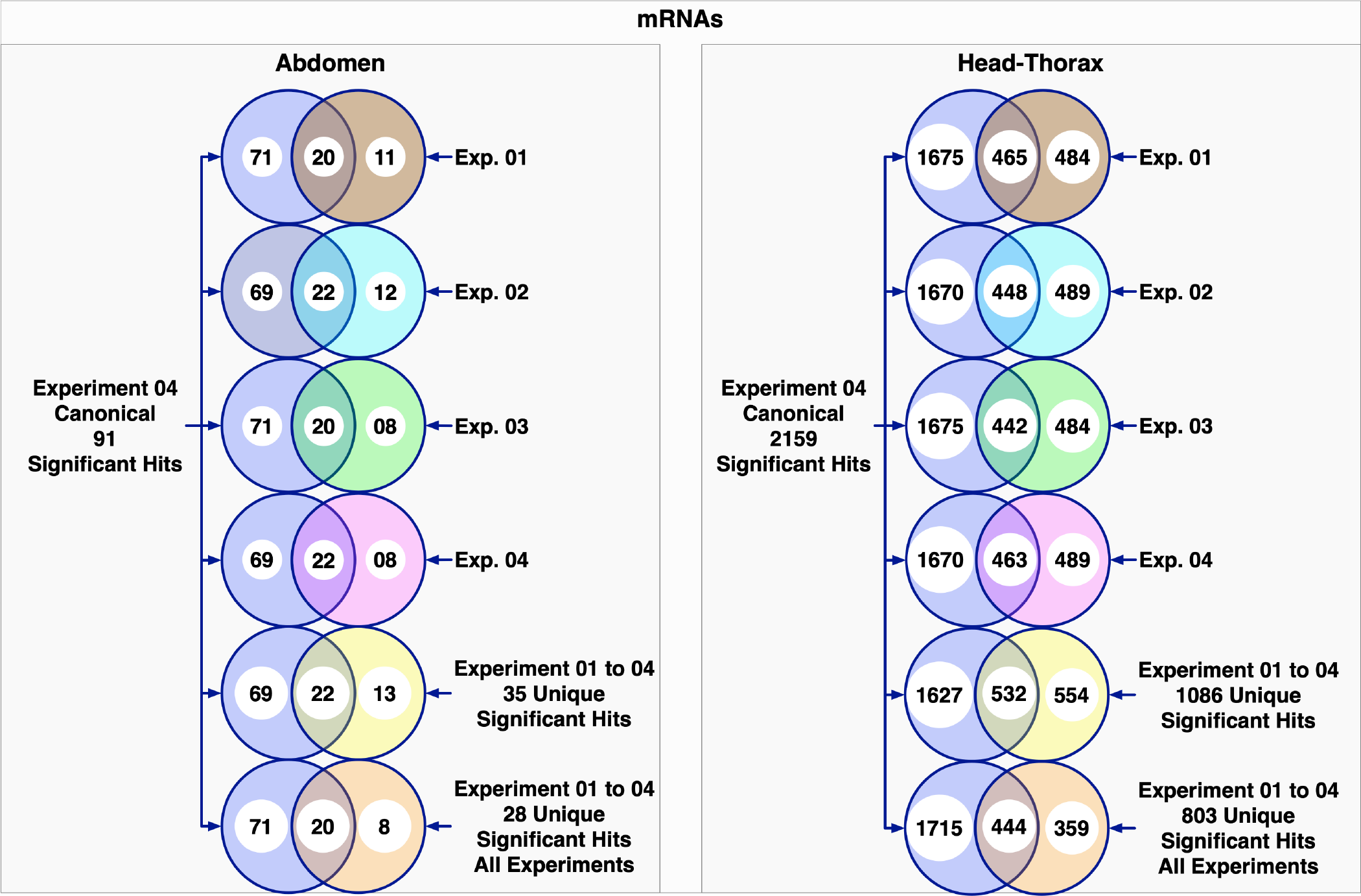
**Using canonical transcripts increased the number of significant hits**. As in Figure 6, here we use Venn diagrams to compare the number of significant genes identified by the Exp. 4 canonical experiment with those identified by Experiments 1 to 4. For Ab, the Canonical experiment (represented in blue) found 91 significantly DE transcripts, 20 of which were transcripts also found by the All experiment. For HT, the Canonical experiment (represented in blue) identified 2,159 significant hits, 444 of which overlapped with the All experiment’s findings.

The majority of the new information was detected by the only canonical transcript experiment (Table 2), demonstrating the profound effect that changing the read assignment addresses has on the overall transcriptome quantification.

#### The Use of Isoforms Versus Canonical Transcripts Determines the Degree of Overlap Of The Number of Differentially Expressed mRNAs (DE-mRNAs) Common to This Study and the 2019 Study

To compare the degree of overlap between our study and Fowler et al. (2019), we used the list of significantly regulated transcripts to extract a list of their corresponding gene IDs (see Materials and Methods).

For Ab, we found that the 28 unique hits detected by the All experiment (Table 1) represent a total of 27 genes (see Materials and Methods and Table 2). Similarly, for HT, we found that the 803 unique hits detected by the All experiment (Table 1) correspond to 753 genes (Table 2). The number of DE genes detected by this and the 2019 study (i.e.overlap) is *∼*17%, for Ab, and *∼*27% for HT (Table 3). Taking both studies’ DE hits into account, the 2019 study detected the majority of unique genes (Fowler et al., 2019) (i.e. 79.8% and 65.8% for Ab and HT, respectively). However, despite our experimental conditions being more stringent, we were able to detect four Ab genes and 154 HT genes that had not been previously detected (Table 3). Of the 27 DE genes in the abdomen, only 23 overlapped with those found in Fowler et al. (2019) (Table 3). We identified 753 genes as being DE in HT, with 599 of those genes found in common with the previous analysis (Table 3).

In the Canonical experiment, we found 91 DE genes in Ab and 2,159 in HT (Table 2). When compared to the genes detected by Fowler et al. (2019), the overlap increased to 54.5% and 57.7% for Ab and HT, respectively (Table 3). A similar increase was also seen for the number of genes detected by only this study when using canonical transcripts (Table 3).

Together, these observations raise serious concerns about the robustness of differential gene expression analysis and our ability to replicate and reproduce any study and should be taken into consideration when comparing different studies.

### Assessment of Individual Differentially Expressed Genes

We selected a set of significant genes only detected by this study and evaluated their potential significance in the PMR. Of the 11 DE genes chosen, six were detected by both the All and Canonical experiments, four were detected by the Canonical experiment alone, and one was detected only by the All experiment. To examine the differing behavior of the All and Canonical experiments, we selected genes coding for one isoform (*UQCR-14*, *CG9498*, *BomS6*, *CG16978*, and *CG43673*), two isoforms (*Epg5* and *CG4267*), three isoforms (*tsl* and *CG8475*), four isoforms (*UGP*), and 12 isoforms (*14-3-3zeta*).

#### Significant Genes Coding for a Single Isoform UQCR-14 (FBgn0030733)

*UQCR-14* codes for a small protein (Q9VXI6, 111aar) that is predicted to be part of the mitochondrial respiratory chain complex III. Interestingly, RNAi of *CG17856*, another gene within this complex, results in increased longevity and resistance to oxidative stress, but also causes a decrease in fertility (Copeland et al., 2009). Reduced lifespan is observed in mated females (Barnes et al., 2008). In both the All and Canonical experiments, this gene’s only transcript, FBtr0074216*^C^*, is significantly upregulated in HT after mating, suggesting it may be involved in the link between lifespan and reproduction (Fig. 8).

**Figure 8.**
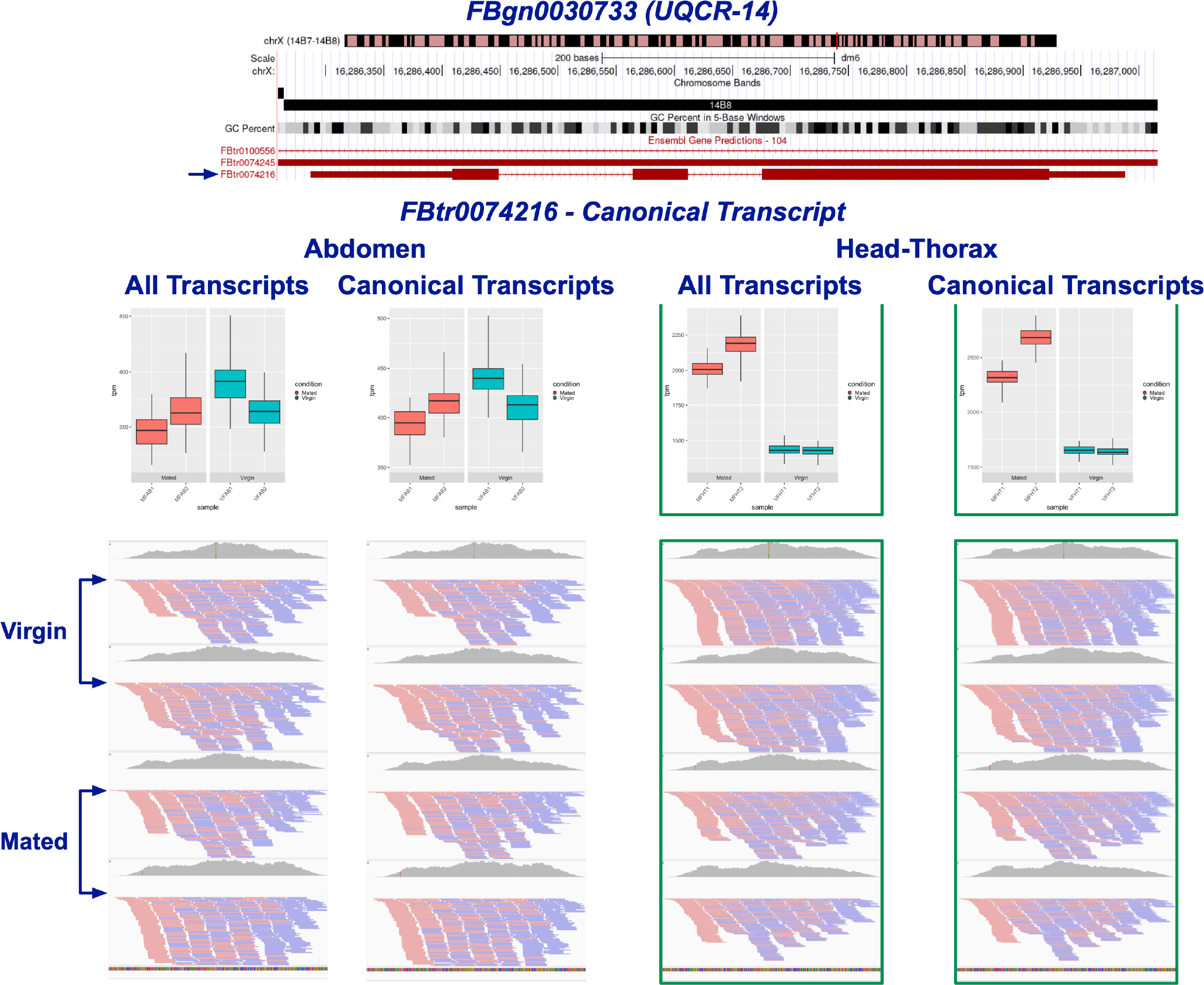
***UQCR-14***. The sole canonical transcript, FBtr0074216*^C^*, is significantly DE in HT after mating in both the All and Canonical experiments (denoted by green outline).

#### CG9498 (FBgn0031801)

*CG9498*, upregulated in Ab after mating, is predicted to be a choline kinase (Q9VMF0, 424aar) with potential chemosensory activity (Shiao et al., 2015) and is associated with climbing ability (Watanabe and Riddle, 2021). Given its upregulation after mating, it is perhaps involved in an enhanced locomotor activity associated with increased foraging and finding an appropriate oviposition site. Interestingly, CG9498 protein was found to interact with Allostatin C receptor 1 (ASTC-R1) (Giot et al., 2003), which is involved in circadian regulation of oogenesis (Zhang et al., 2021a). The gene’s sole transcript, FBtr0079282*^C^*, was detected in both the All and Canonical experiments (Fig. 9).

**Figure 9.**
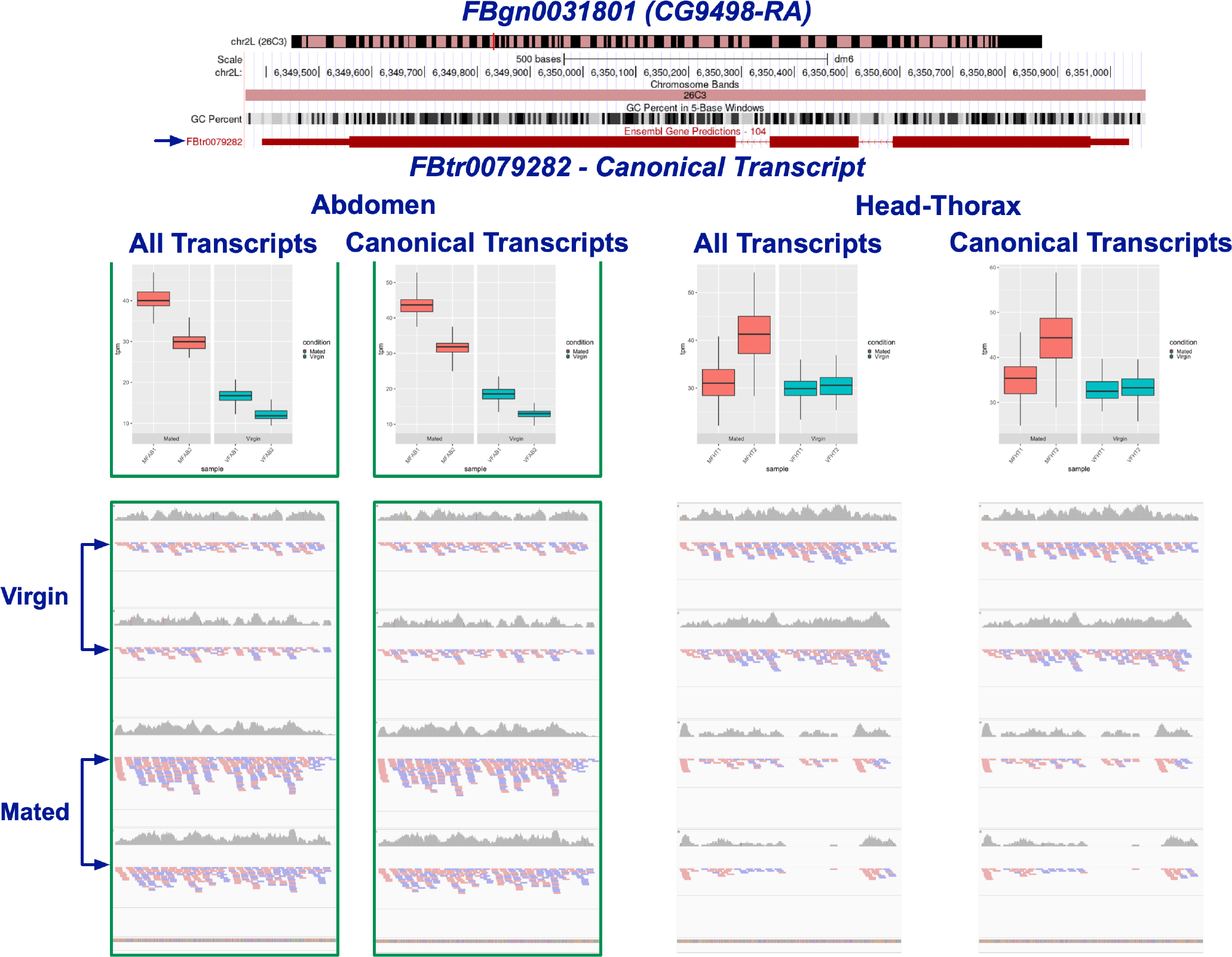
***CG9498***. The sole canonical transcript, FBtr0079282*^C^* is upregulated in Ab after mating in both the All and Canonical experiments (denoted by green outline).

#### BomS6 (FBgn0040733)

*BomS6* is a antimicrobial peptide-like gene induced by Toll signaling (Lin et al., 2019). Its transcript FBtr0086727*^C^*, coding for a small secreted peptide (A1ZB64, 40 aar) was found to be significantly upregulated in HT after mating in the Canonical experiment (Fig. 10).

**Figure 10.**
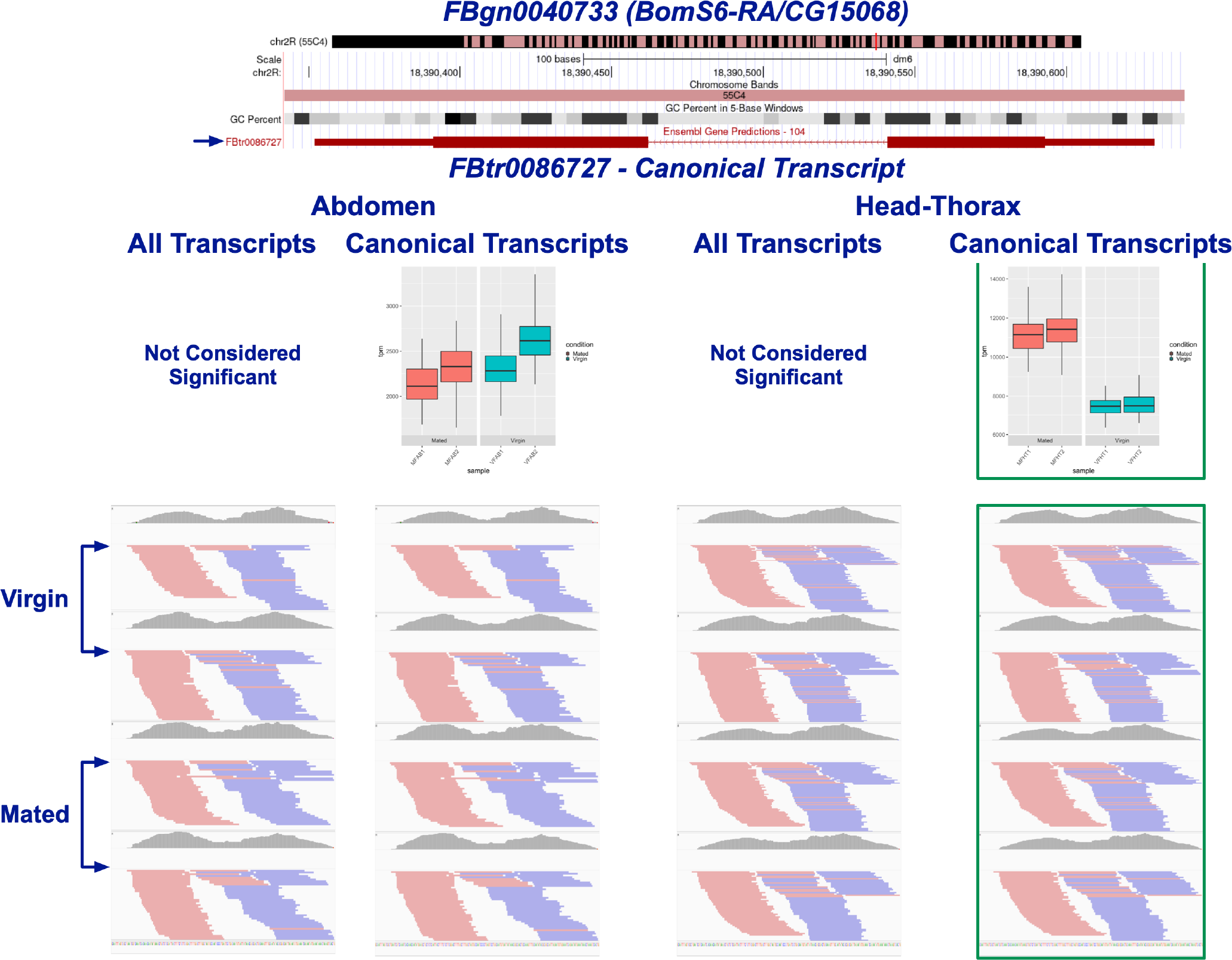
***BomS6***. The sole canonical transcript, FBtr0086727*^C^* is upregulated in HT after mating in the Canonical experiment only (denoted by green outline).

#### CG16978 (FBgn0040972)

*CG16978* codes for an uncharacterized peptide (Q9VK38, 96 aar), but is predicted to be involved in immune function, perhaps serving an antimicrobial function (De Gregorio et al., 2001, 2002). Transcript FBtr0307212*^C^* was detected by both the All and Canonical experiments as being upregulated in HT after mating (Fig. 11).

**Figure 11.**
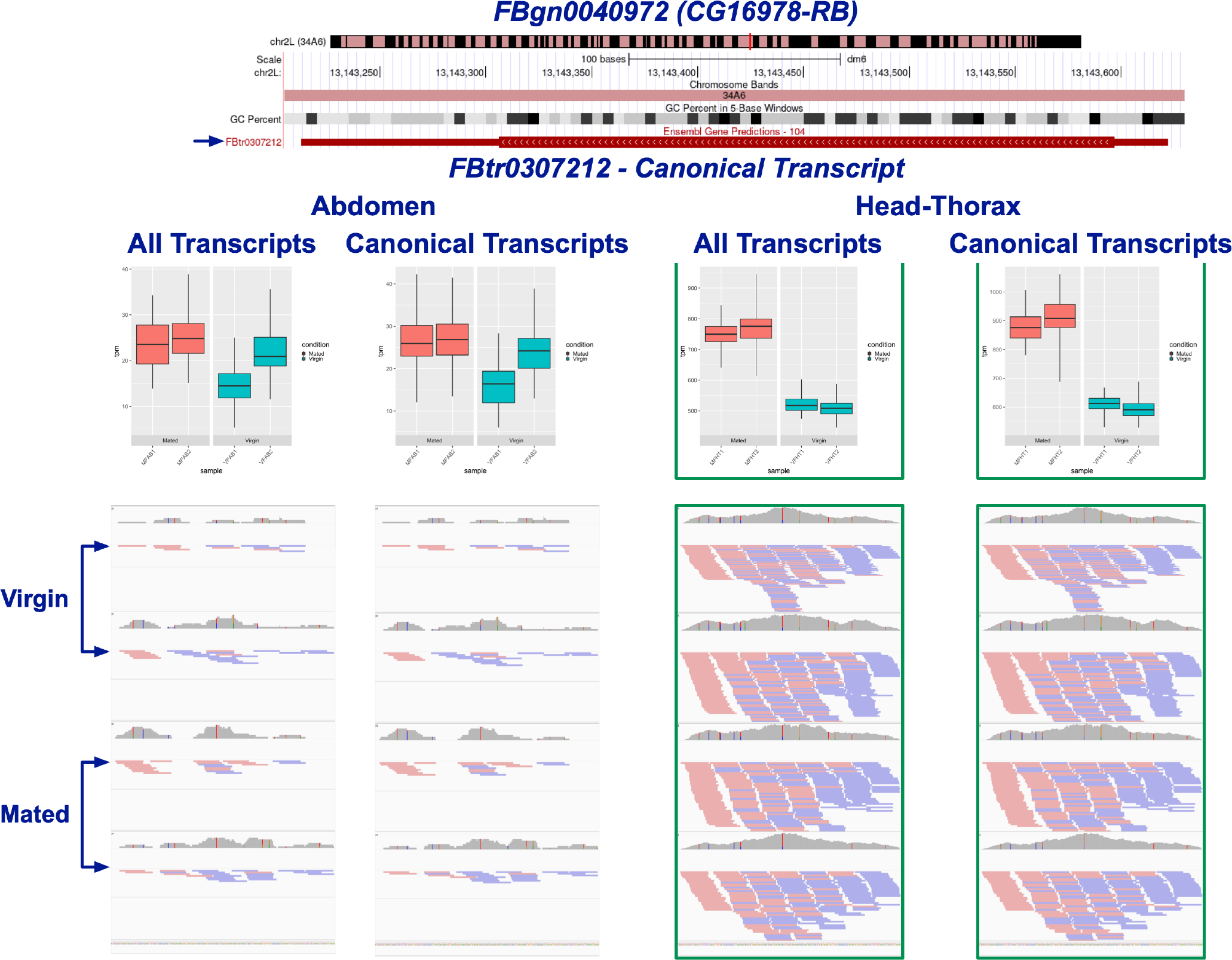
***CG16978***. The sole canonical transcript, FBtr0307212*^C^* is upregulated in HT after mating in Both the All and Canonical experiments (denoted by green outline).

#### CG43673 (FBgn0263748)

*CG43673* is uncharacterized, but predicted to code for a 230 aar extracellular protein involved in chitin binding (M9PEP7). Its transcript, FBtr0310395*^C^*, was detected to be upregulated in HT after mating by both the All and Canonical experiments (Fig. 12).

**Figure 12.**
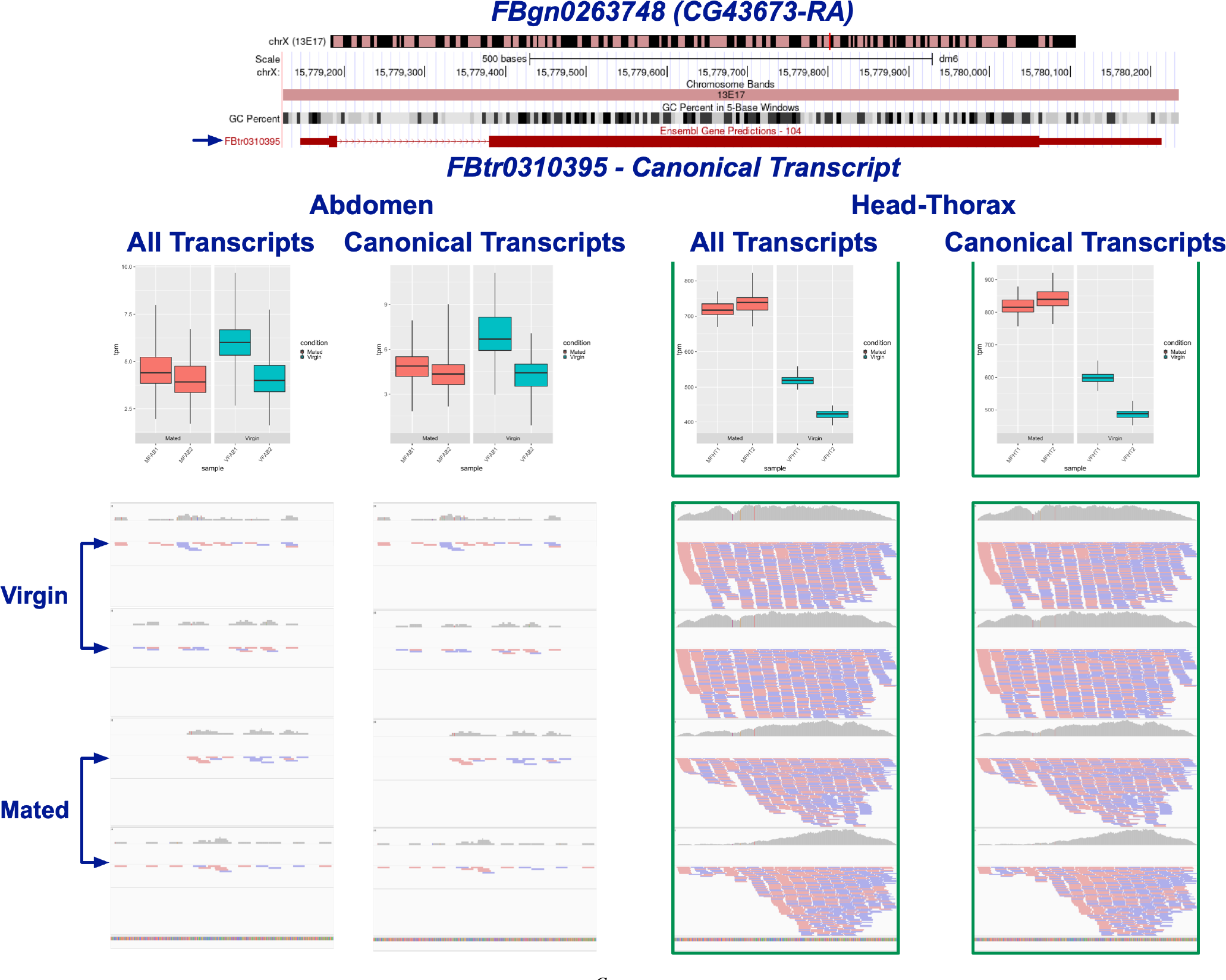
***CG43673***. The sole canonical transcript, FBtr0310395*^C^* is upregulated in HT after mating in the All and Canonical experiments (denoted by green outline).

#### Significant Genes Coding for Two Isoforms Epg5 (FBgn0038651) - An Enigma

*Epg5*, codes for two transcript isoforms, FBtr0273382*^C^* and FBtr0305051, both predicted to encode nearly identical proteins involved in the later stages of the autophagy pathway. FBtr0273382*^C^* codes for a 2455 aar protein (Q9VE34), while isoform FBtr0305051 codes for a 2454 aar protein (A0A0B4K6N7). The only difference between these two protein isoforms is the presence of an extra Lysine (K) residue at position 11 of Q9VE34. Both transcript isoforms were declared to be significant by the All experiment, though in opposite directions. We observed that FBtr0273382*^C^* was repressed in HT after mating (Fig. 13), while FBtr0305051, which showed no expression in abdomen, was declared to be upregulated in HT (Fig. 14). These observations are both puzzling and contradictory. Although they initially suggest the presence of exciting isoform-specific regulation, we cannot accept these results without intensive extra experimental evidence. Visual inspection of the read coverage distribution that mapped to each transcript reveals few differences consistent with the isoform-specific regulation declared by these pipelines (Figs. 13 and 14). These quasi-identical transcripts should have been similarly detected in all conditions. Identical transcripts are removed during Salmon index construction, and inspection of our log files confirmed that neither transcript of *Epg5* was affected by this behavior of Salmon.

**Figure 13.**
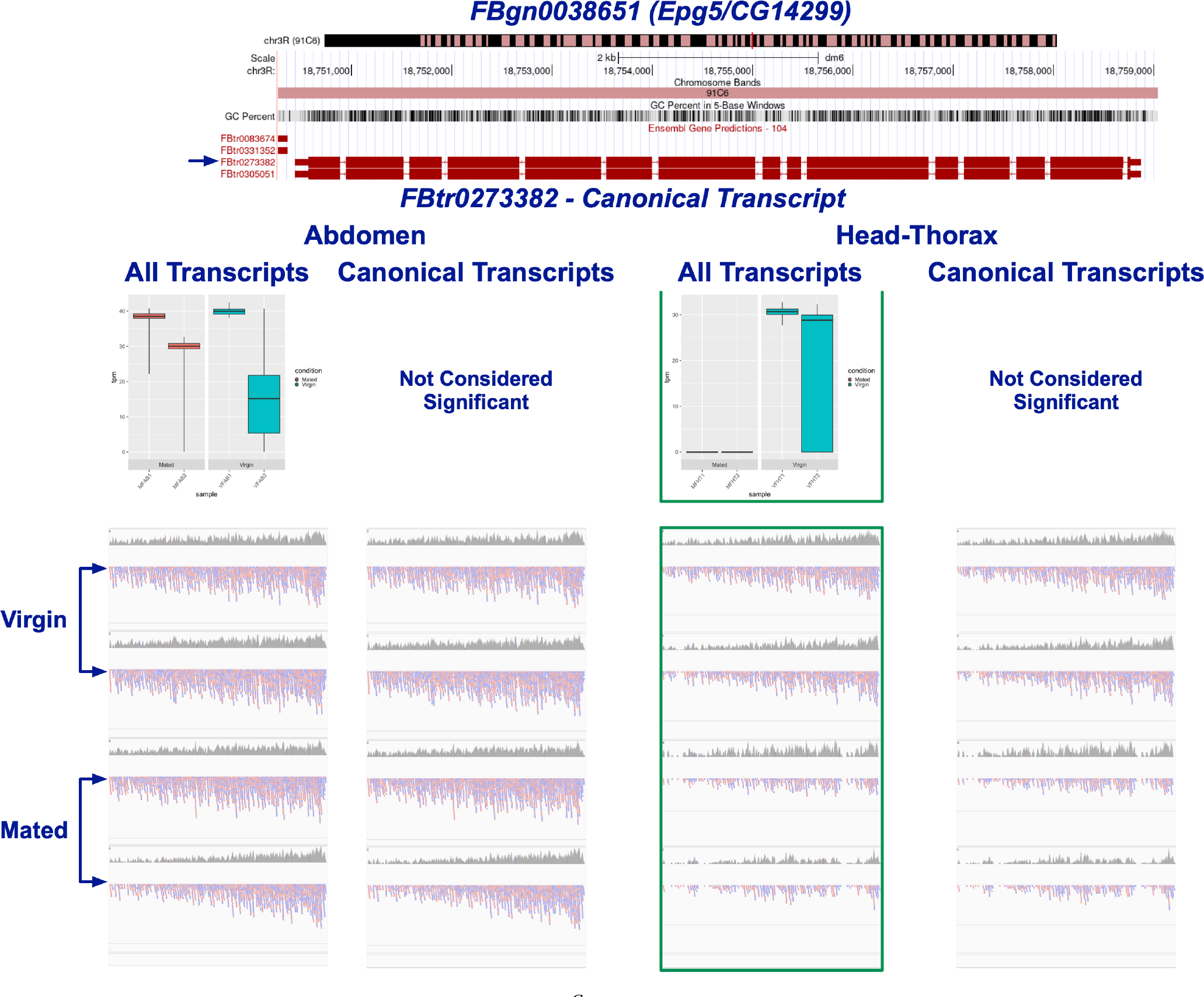
***Epg5* canonical transcript**. Transcript FBtr0273382*^C^*is downregulated in HT after mating in the All experiment only (denoted by green outline).

**Figure 14.**
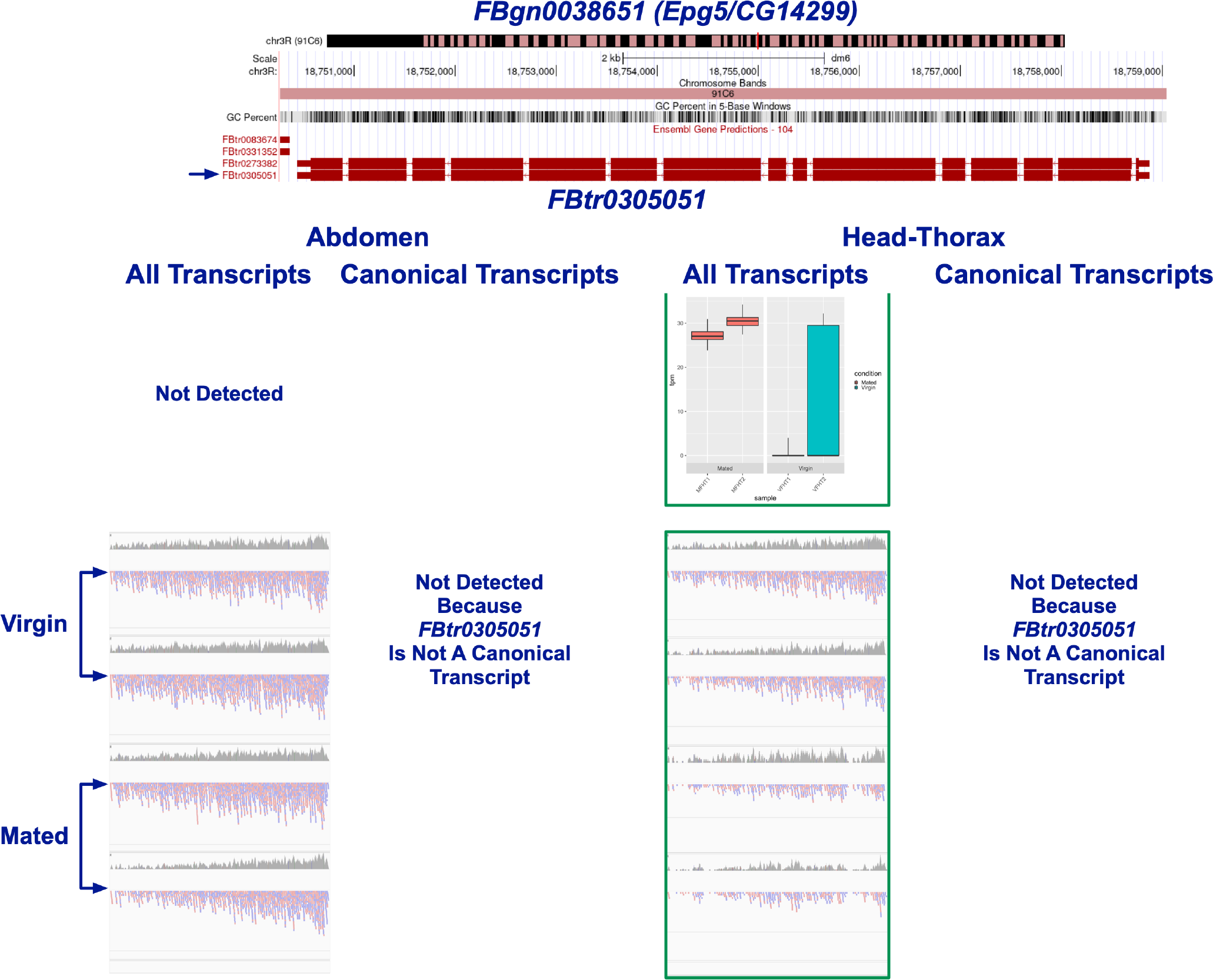
***Epg5* alternative transcript isoform**. Transcript FBtr0305051 was declared to be upregulated in HT after mating in the All experiment (denoted by green outline).

Interestingly, *Epg5*-specific defects in autophagy are directly linked to reduced male courtship and reproductive success in zebrafish (Fontana et al., 2021). When autophagy is interrupted in male Drosophila, mating behavior defects are observed, perhaps due to altered synaptic regulation impairing olfactory perception (Ratliff et al., 2015). Given the importance of olfaction and other chemosensory input for PMR changes in receptivity and food sources (Hussain et al., 2016; Zhou et al., 2014), it is possible that Epg5 has a role in autophagic pathways involved in PMR synaptic plasticity.

#### CG4267 (FBgn0264979)

*CG4267* was significantly DE in Ab after mating in both the All and Canonical experiments. This gene codes for two transcript isoforms, FBtr0335159*^C^*and FBtr0077751 (Figs. 15 and 16). Interestingly, the canonical transcript is upregulated after mating and the alternative isoform is downregulated. While both isoforms code for the same secreted lipase protein (Q9VQ96, 374aar), FBtr0335159*^C^*has a significantly longer 3’-UTR (Fig. 15), which is likely to be involved in regulating its own expression.

**Figure 15.**
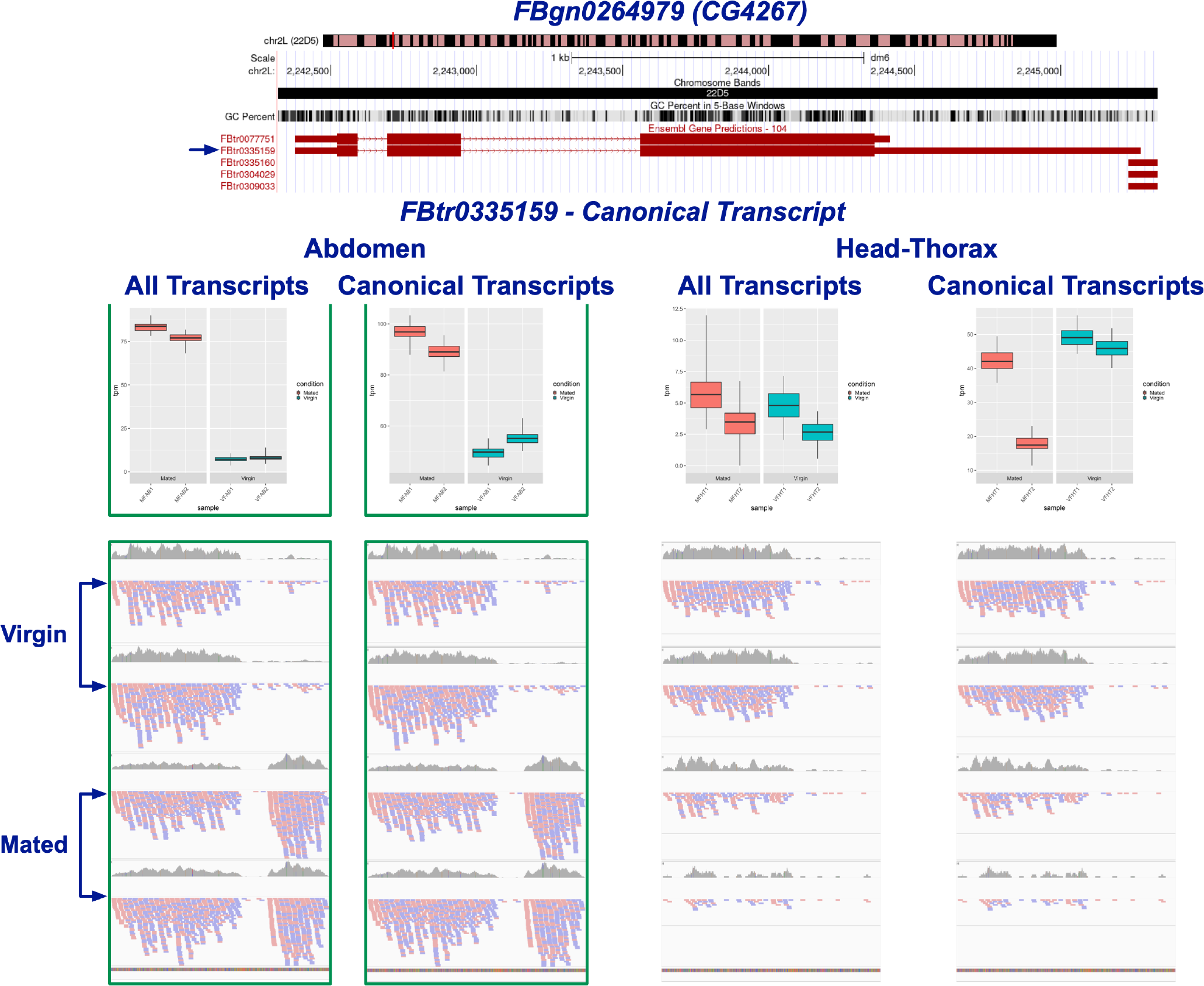
***CG4267* canonical transcript**. Transcript FBtr0335159*^C^* is upregulated in Ab after mating in both the All and Canonical experiments (denoted by green outline). Note the prominent read coverage in the 3’-UTR region of the Ab mated condition for the All and Canonical experiments. We believe this area may represent a different, unannotated transcript that is causing this transcript to be called as upregulated.

**Figure 16.**
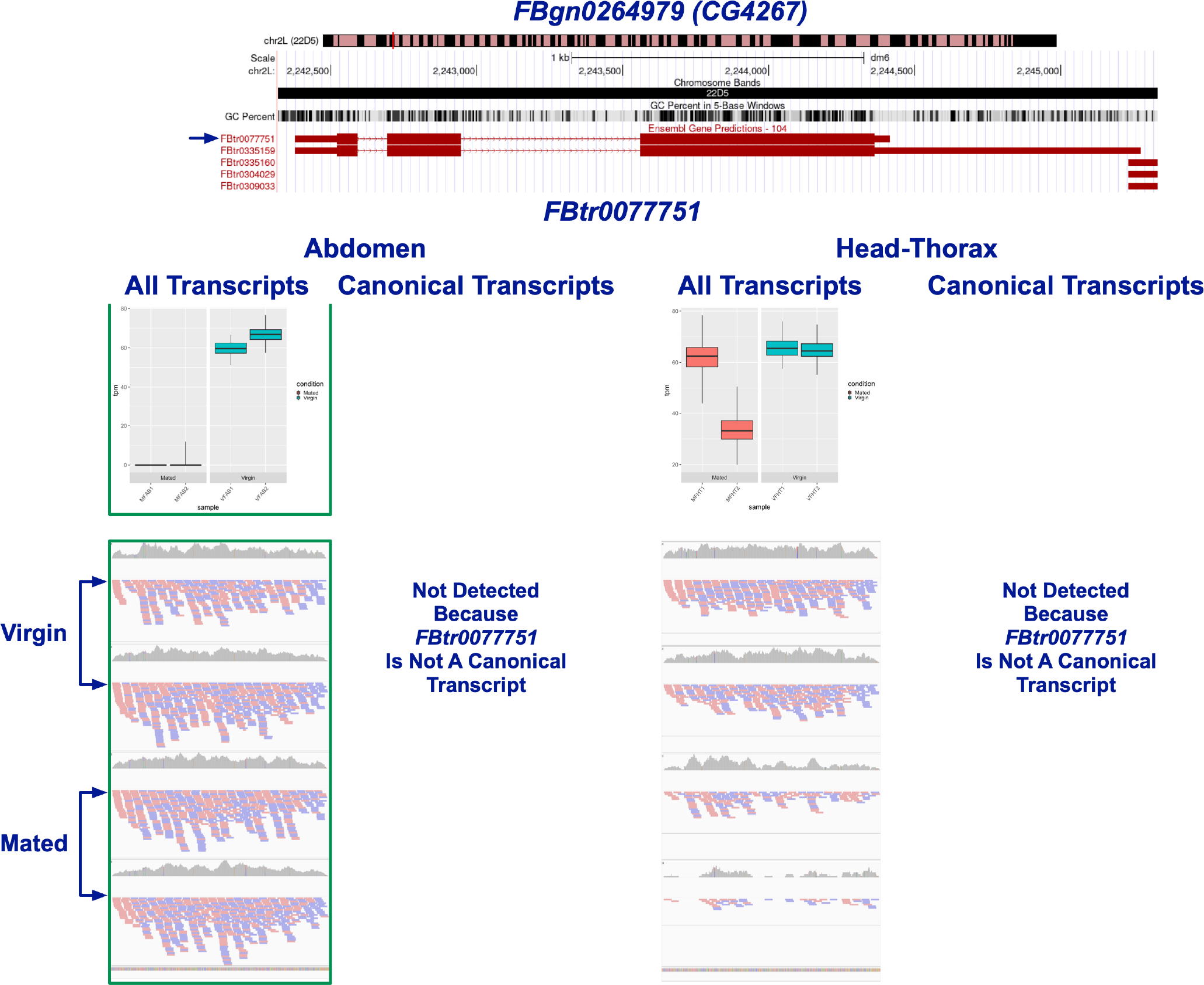
***CG4267* alternative transcript isoform**. Transcript FBtr0077751 is considered downregulated after mating by the All experiment in Ab (denoted by green outline).

*CG4267* is a predicted lipase and has been identified as part of a syntenic block of genes flanking rapidly evolving serine proteases implicated in sexual conflict (Lawniczak and Begun, 2007). Additionally, *CG4267* is upregulated in S2 cells early after infection with *Buchnera aphidicola*, indicating a role in immune response (Douglas et al., 2011). Taken together, these previous studies suggest that *CG4267* is involved in mitigating female harm from mating by processing male reproductive molecules and responding to infection. Though FBtr0335159*^C^* being upregulated (Fig. 15) and FBtr0077751 being downregulated in the mated condition (Fig. 16) seems puzzling, we suspect is likely to be an artifact of the lack of experimental replicates. The read coverage distribution observed for transcript FBtr0335159*^C^*, although predominant in the region corresponding to the 3’-UTR in the mated condition, does not display an expected continuous coverage with the main part of the transcript (Fig. 15), skewing the results to show an increase in expression post-mating. This read distribution behaves as if those reads were part of their own independent, and obviously unannotated, transcript unit. These observations thus require further experimental verification.

#### Significant Genes Coding for Three Isoforms tsl (FBgn0003867)

*tsl* was found to be upregulated in HT after mating by the Canonical experiment (Fig. 17). This gene codes for three isoforms: FBtr0334312*^C^*, FBtr0084164, and FBtr0084165, all of which are predicted to code for the same 353 aar protein (P40689), a ligand expected to interact with the torso receptor.

**Figure 17.**
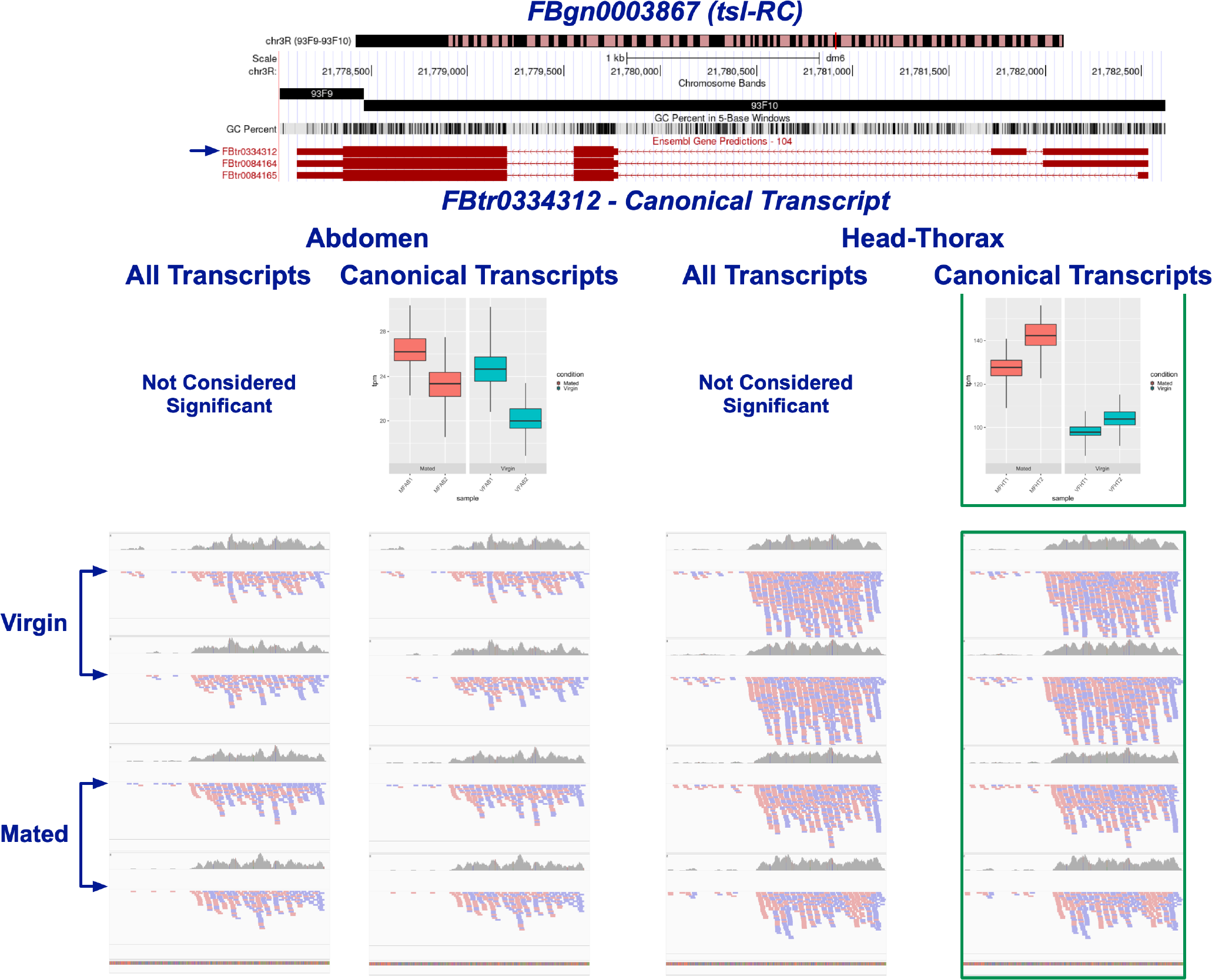
***tsl* canonical transcript**. Transcript FBtr0334312*^C^* was found to be upregulated in HT after mating by the Canonical experiment (denoted by green outline). The biased read distribution towards the 3’-end of the gene model suggests that the reads assigned to this transcript in the Canonical experiment actually correspond to one of the other two *tsl* isoforms (see Figs. S3 and S4).

While *tsl* is required during oogenesis for patterning in early embryos (Savant-Bhonsale and Montell, 1993), its significant expression in HT of mated females implies alternative functions. *tsl* mutants display growth defects mirroring those seen when insulin signaling is disrupted and is purported to act within the insulin signaling pathway (Henstridge et al., 2018). Insulin signaling affects both egg laying and remating latency in mated flies (Wigby et al., 2011; Yang et al., 2008). Additionally, the pathway triggers the production of hormones related to PMR: juvenile hormone, which in turn stimulates the production of the steroid prohormone ecdysone in the ovaries (Tu et al., 2002, 2005). Investigating the role of mating- induced *tsl* expression, as it relates to insulin signaling, will provide a more nuanced understanding of the factors influencing PMR.

An examination of the read coverage distribution to these different transcript isoforms raises some issues. First, clearly, while FBtr0334312*^C^* is upregulated, the bulk of the reads do not cover the entirety of the transcript, with the 5’ end showing significantly lower coverage (Fig. 17). In contrast, though the remaining two transcripts were not declared as significant, the read coverage distribution of FBtr0084165 is uniform and covers the entirety of its transcript (see Figs. S3 and S4). While we believe *tsl* is significantly DE after mating, we suspect that the presence of other isoforms confounded the identification of this gene as significant in the All transcripts experiments. In addition, we think that the majority of the quantified reads of the Canonical experiment might in fact correspond to the FBtr0084165 transcript.

#### CG8475 (FBgn0031995)

*CG8475* codes for three transcript isoforms: FBtr0335186*^C^*, FBtr0079638, and FBtr0079639. FBtr0335186*^C^*codes for a 1098 aar protein (M9PCI5), FBtr0079638 codes for a 1093 aar protein (Q9VLS1), and FBtr0079639 is predicted to encode a 506 aar protein. Though FBtr0335186*^C^* was declared as significant only in the Canonical experiment, it should be noted that this transcript was also found to be significant in Exps. 3 and 4, but not in Exps. 1 and 2, preventing it from meeting the threshold of significance in the All experiment. While our observations are consistent with *CG8475* being significantly upregulated in HT after mating, it looks to us that the read coverage distribution of all three predicted isoforms is not uniform along the predicted canonical isoform transcript (see Fig. 18, S5, and S6). This suggests the strong possibility that the observed transcript read coverage is not consistent with the existing annotated gene models.

**Figure 18.**
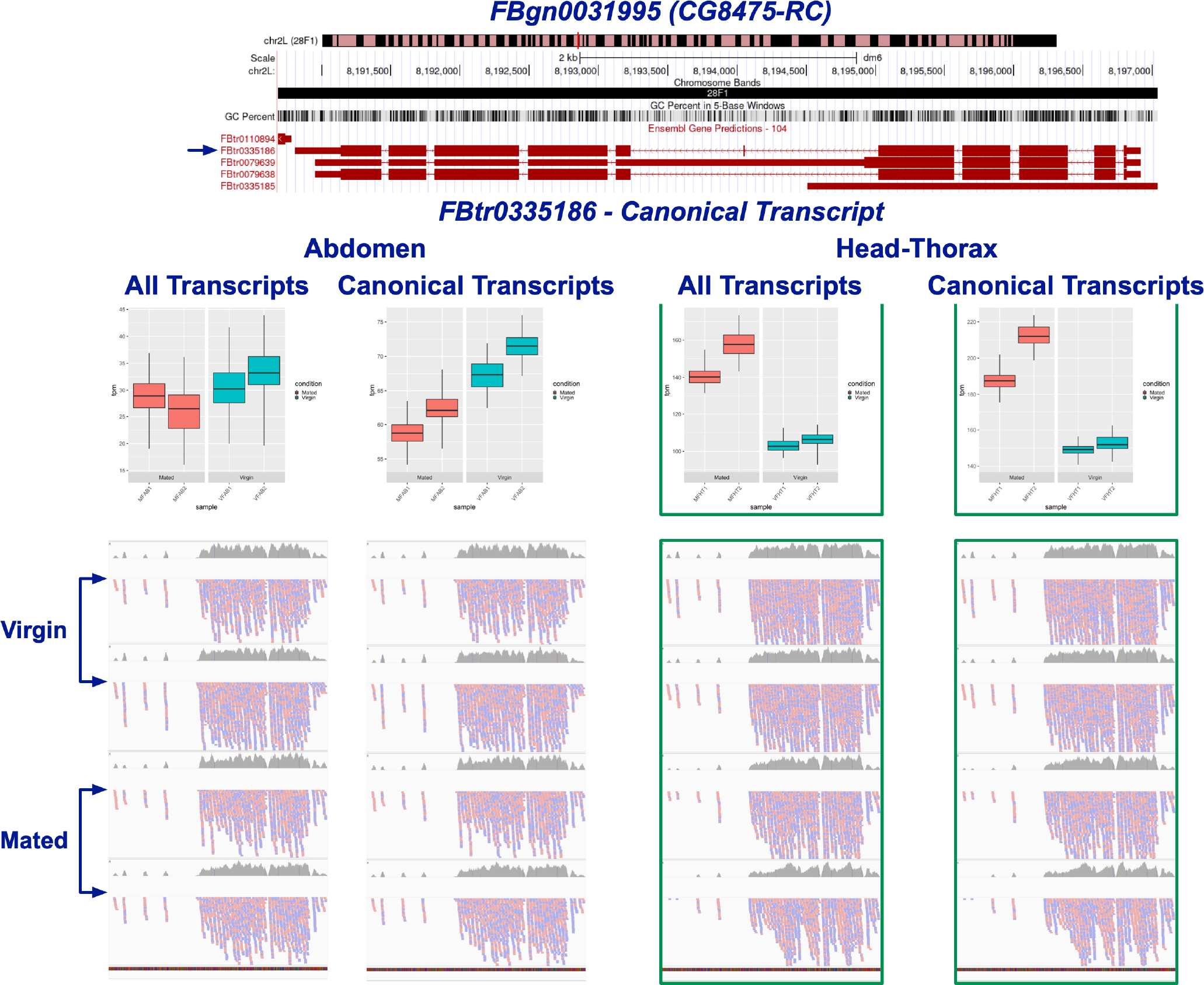
***CG8475* canonical transcript**. Transcript FBtr0335186*^C^*was found to be significantly upregulated in HT after mating (denoted by green outline) by the All and Canonical experiments. Note that the read distribution is not even over the length of the transcript, suggesting that the transcript observed in this study is not consistent with the current gene model.

A GWAS study identified *CG8475* as being associated with midgut mitosis, and its downregulation in enteroendocrine cells (EEs) reduces the division of intestinal stem cells (Tamamouna et al., 2020). Mating is known to induce cell proliferation in the gut via juvenile hormone signaling (Reiff et al., 2015). Ecdysone, which is produced in the ovaries after mating, also increases proliferation of gut stem cells (Ahmed et al., 2020). Interestingly, a factor released from EEs in the midgut has been shown to induce proliferation of germline stem cells after mating (Ameku et al., 2018). Cross-talk among organs and tissues to coordinate cellular responses to mating is an important factor in PMR. Given the post-mating upregulation of *CG8475* seen in our data, perhaps it is a member of the complex pathways increasing gut cell proliferation in mated flies.

#### Significant Genes Coding for Four or More Isoforms UGP (FBgn0035978)

*UGP* codes for four transcript isoforms. FBtr0076501*^C^* codes for a 520 aar protein (Q9VSW1) and FBtr0301699, FBtr0076500, and FBtr0301698 code for smaller 518, 513, and 511 aar proteins, respec- tively. The product of the canonical isoform is an enzyme involved in carbohydrate metabolism. This enzyme works in a pathway also including *jhamt*, an enzyme that processes JH into its active form (Schlamp et al., 2021).

This gene was found to be significantly upregulated after mating in HT by the Canonical experiment (see Figs. 19 and S7-S9). Note that not all predicted isoforms have the expected read coverage distribution (see Fig. S7). This suggests that, in the case of FBtr0301699, which happens to be the largest predicted transcript for this gene (2,686 bp versus 2,102 bp for the canonical transcript), the reads that mapped to this transcript do not necessarily come from this transcript. This observation underscored, yet again, the potential for read “cross-contamination”, or misassignment, that can seriously compromise the evaluation of transcriptional regulation.

**Figure 19.**
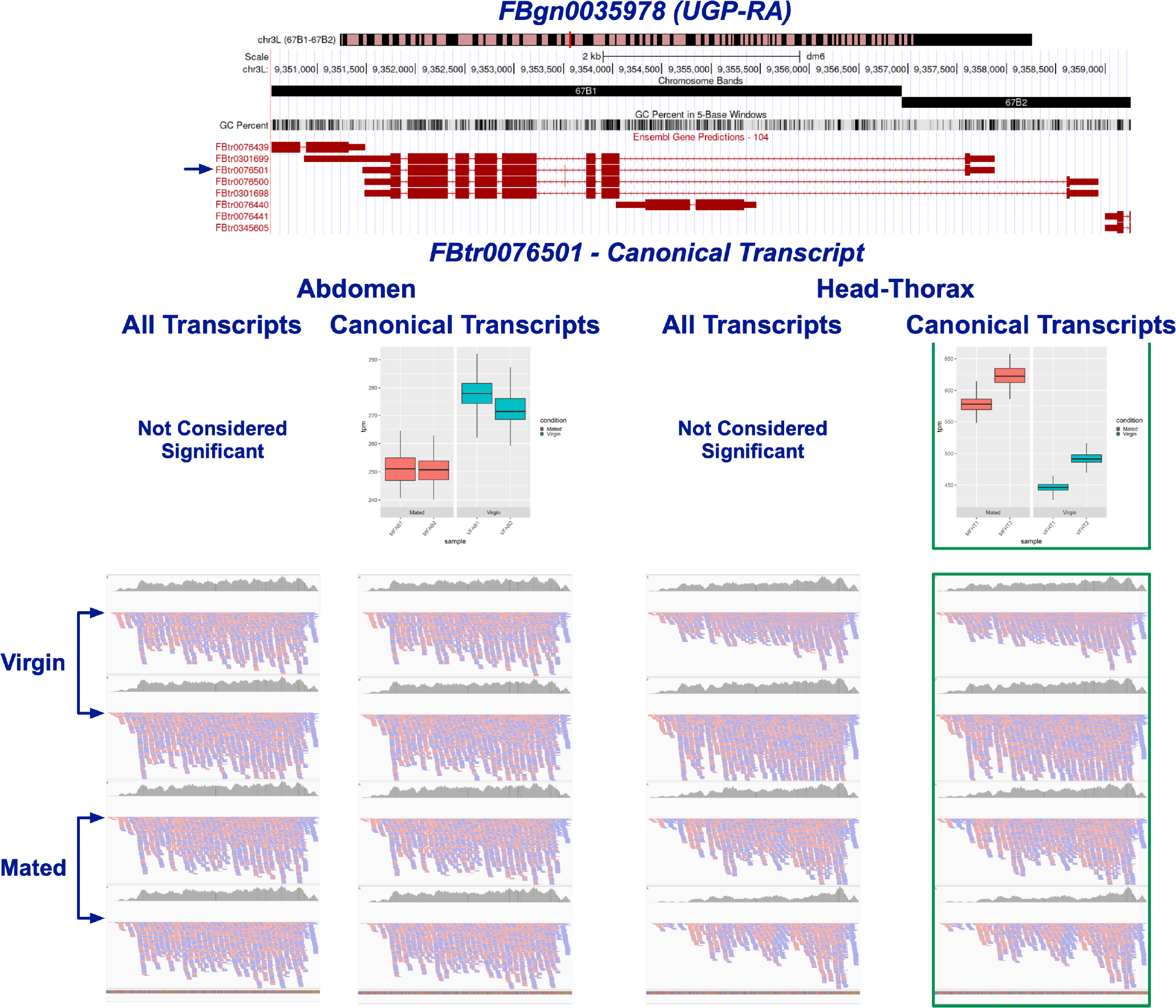
***UGP* canonical transcript**. Transcript FBtr0076501*^C^*was found to be significantly upregulated in HT after mating in the Canonical experiment (denoted by green outline).

#### 14-3-3zeta (FBgn0004907)

*14-3-3zeta* represents a beautiful example of complexity, in that the gene is predicted to encode for a total of 12 transcript isoforms, all of which code for the same 248 aar protein (P29310). While this gene was found to be significant by both the All and Canonical experiments, in each case, different transcript isoforms were considered significant (Figs. 20 and S10-S20). For example, FBtr0088415 and FBtr0088416 were declared significant by experiments 1-4 (All). In contrast, FBtr0100182 was declared significant by experiments 1, 3, and 4 and FBtr0100183 was significant in experiments 1 and 3.

**Figure 20.**
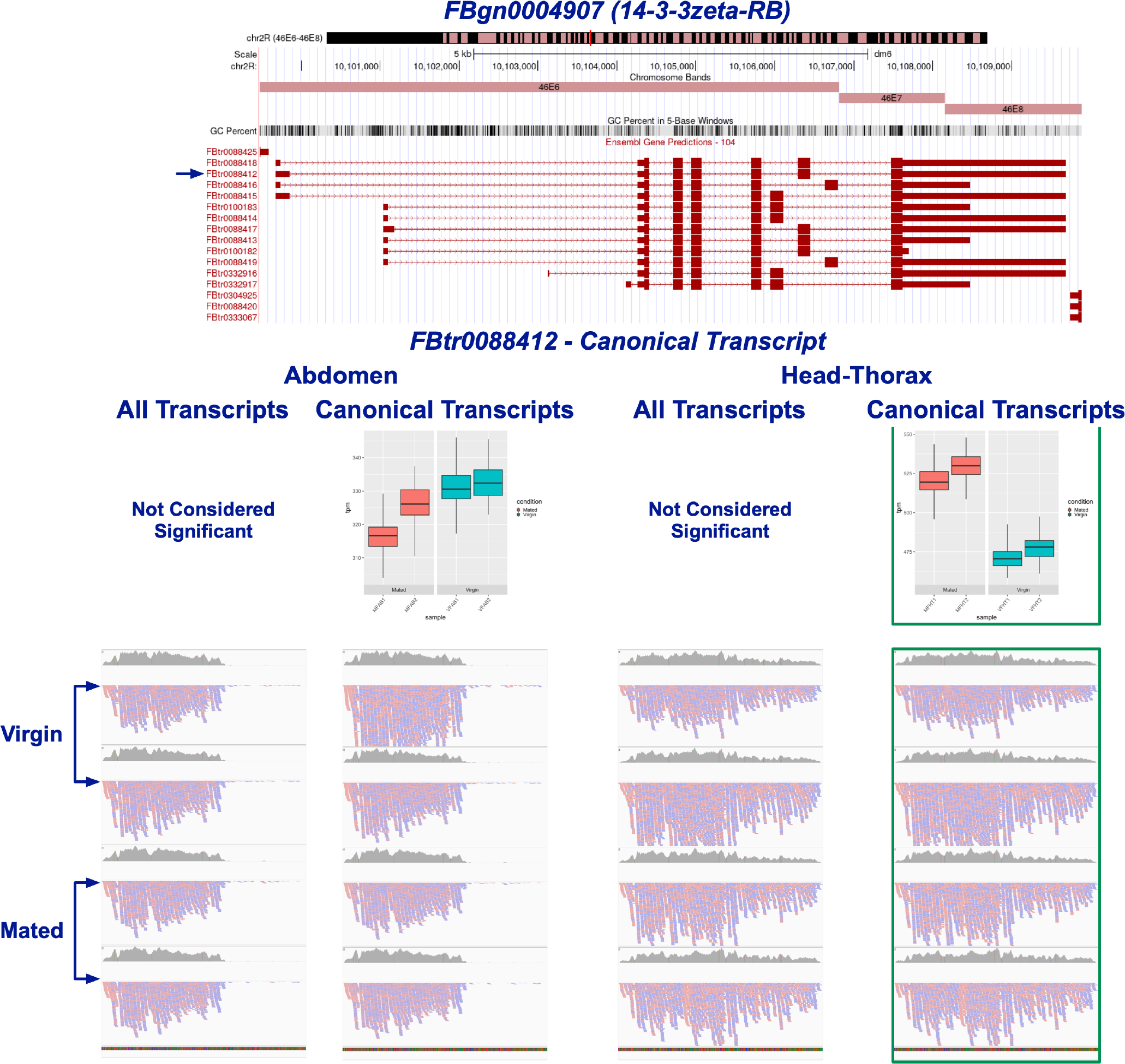
***14-3-3zeta*/*leo* canonical transcript**. Transcript FBtr0088412*^C^*, one of 12 transcripts for this gene, was found to be significantly DE in HT by the Canonical experiment (denoted by green outline). Four of the other 11 transcripts were also found to be significant in the All experiment (see Figs. S10-S20).

While *14-3-3zeta* is clearly upregulated in HT after mating, these results underscore the read-to-isoform assignment problem and how this affects the evaluation of isoform transcriptional activity. Assuming that all the predicted isoforms are real and have a defined biological function, given that all of them are predicted to encode the same protein, suggests that their existence is likely to be related to the presence of regulatory mechanisms operating in an isoform-specific manner. In other words, different transcript isoforms must have sequences that are the target of a regulatory mechanism controlling their expression and/or stability, thus indirectly regulating the presence of the gene’s protein product. Our results, again, highlight the need to evaluate gene expression taking into consideration the existence of isoform complexity and how they potentially act as “sponges” to confuse the evaluation of transcriptional significance.

It is nevertheless exciting to implicate *14-3-3zeta* in PMR biology. This gene, also known as *leonardo* (*leo*), is expressed in the mushroom bodies (MB). *leo* is involved in associative learning and olfactory memory (Skoulakis and Davis, 1996). The MB plays an important part in oviposition and laying site selection behavior after mating (Azanchi et al., 2013; Fleischmann et al., 2001). Mated females prefer to lay eggs in substrate where they have observed others lay (Battesti et al., 2012), where they also use the presence of the pheromone 11-cis-vaccenyl acetate deposited by other females to confirm site selection (Dumenil et al., 2016). *leo* may be involved in integrating the social learning and olfactory perception aspects of egg laying in mated females.

### Differential Non-Coding RNA (ncRNA) Expression Analysis

This study is fundamentally different from that of Fowler et al. (2019), in that we determined the differential regulation of all ncRNAs, not just miRNAs. Unlike what we observed for the mRNAs, only one gene, *miR-927* (FBgn0262210/FBtr0304254), was found to be DE in all four experiments in abdomen, though we did observe differential ncRNA regulation for most of the experiments performed (see Table 1 and Figs. 21 and S2). The vast majority of the ncRNA reads that were processed in these experiments (*∼*80%), failed to align to any of the indexes used (see Fig. 5). The majority of these reads belonged to the bacteriophage *PhiX 174*, a spike-in control used for Illumina sequencing runs. We also observed the presence of reads belonging to unannotated *Drosophila* insertion elements.

**Figure 21.**
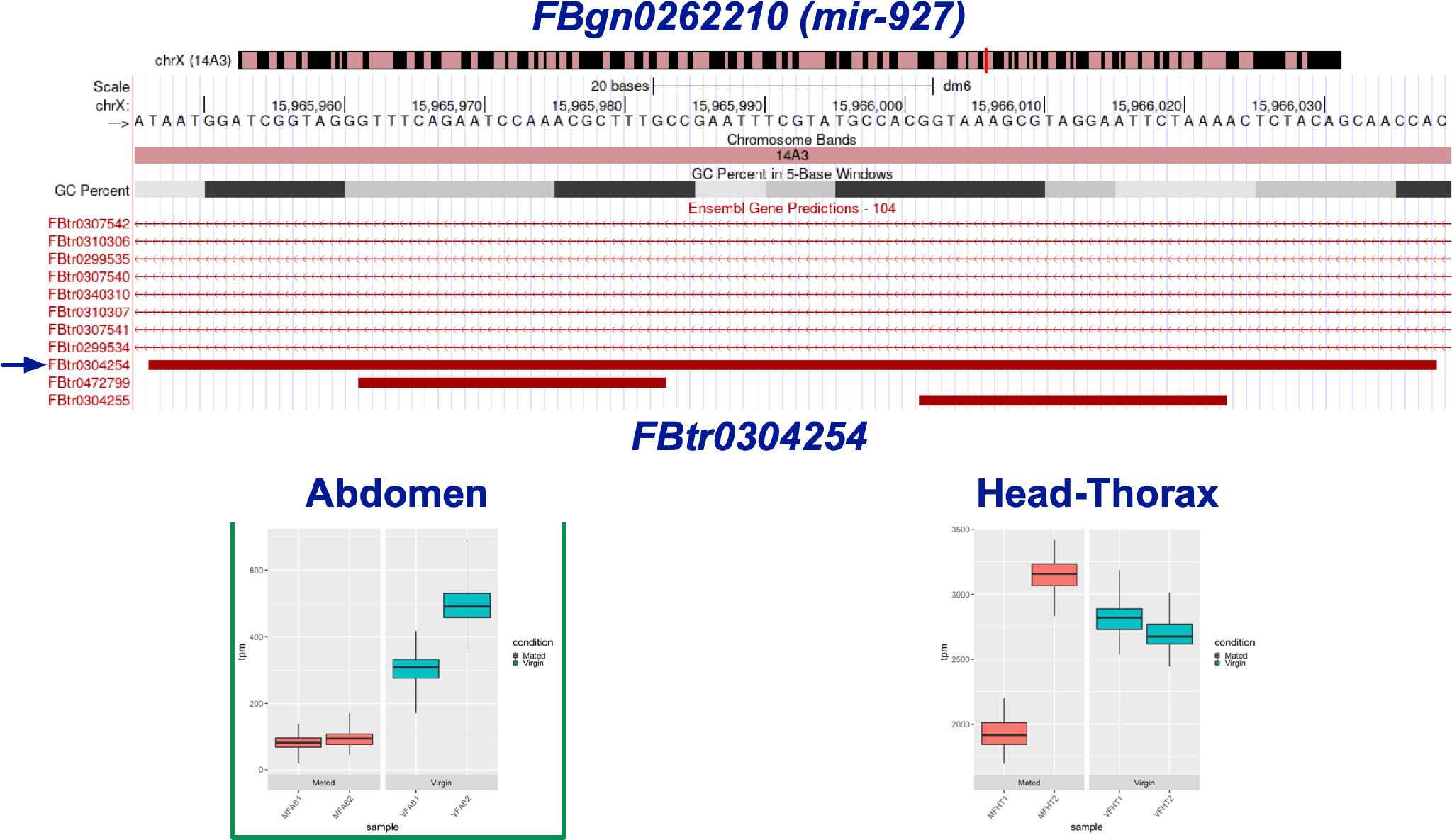
***miR-927***. *miR-927* was the only ncRNA found to be DE in the All experiment. It is downregulated in Ab after mating.

*miR-927* was found to be significantly downregulated in Ab after mating. Given that miRNAs generally downregulate their mRNA targets via 3’-UTR hybridization, we would expect that target genes of *miR-927* may be upregulated after its mating-induced downregulation. Though there are no predicted targets (using TargetScanFly (Agarwal et al., 2015)) of *miR-927* that are significantly DE in this analysis, *miR-927* has been associated with some post-mating effects.

The conserved transcription factor *Kruppel homolog1* (*Kr-h1*) is a demonstrated target of *miR-927*, containing two miR-927 binding sites in its 3’-UTR (He et al., 2020). Overexpression of *miR-927*, with a resulting decrease in Kr-H1 protein, in the fat body of adult females reduces oviposition, a vital component of the PMR. The decrease in *miR-927* after mating may be the result of increased levels of juvenile hormone, which has been shown to target the miRNA (He et al., 2020) and is itself spurred on by the receipt of Sex Peptide (SP) during mating (Moshitzky et al., 1996). Juvenile hormone is required for the maturation of oocytes (Soller et al., 1999), further tying *miR-927* to the female PMR.

## FINAL CONSIDERATIONS AND CONCLUSIONS

### About Replicability, Reproducibility and Experimental Design

Reproducibility in RNA-seq analysis is a complicated issue, with insufficient reporting of methods being a confounding factor. Within a peer-reviewed article, one should expect that all necessary information needed to replicate the experiments, and ideally the results, be included. Given that raw sequencing data is available in repositories and methods are described in the associated papers, RNA-seq results should be simple to replicate. However, it has been found that only one-quarter of RNA-seq papers include enough information to reproduce the experiments, with even fewer including the specific parameters used during analysis that would be necessary for complete replication (Simoneau et al., 2021). When it comes to computational analysis of existing data, two big questions are raised: can the original lab repeat their own results and can a different lab repeat the original team’s results? For the latter question, it is not necessarily a matter of reproducing the experiment exactly, but rather reproducing the interpretations of the data and conclusions made during the original study.

The experimental design process is critical to ensure the statistical power and biological relevance of differential gene expression studies. Although the cost of RNA sequencing has decreased over recent years, most research groups still must make a trade-off when deciding between increasing the sequencing depth of their samples or increasing the number of samples (i.e. replicates) at the expense of some sequencing depth. Library preparation is the major cost associated with additional samples, thus groups may try to gather more reads from fewer samples (Tarazona et al., 2011). Liu et al. (2014) determined that increasing sequencing depth beyond 10M reads has diminishing returns, and that increased biological replicates with less depth is a better approach for DGE studies (Liu et al., 2014). For example, 30M reads are best applied to 10M reads for three samples as opposed to 15M reads for just two samples (Liu et al., 2014). Interestingly, RNA-seq experiments require little in the way of technical replication due to well-controlled sequencing techniques that introduce little to no variability (Marioni et al., 2008), but biological replicates are still necessary (Hansen et al., 2011). Using more biological replicates results in better resolution of low fold change genes between conditions and allows for better conclusions to be drawn about the effect of the studied treatment.

### Conclusions

In this work, we examined if we could both reproduce and improve the detection of significantly-regulated transcripts using an important, previously described dataset. We found that the set of significant genes detected was not only highly dependent on the number of transcripts entering the evaluation (i.e., All versus only Canonical), but also that the number of isoforms associated with each individual gene affected the outcome. The increasing accessibility of next-generation sequencing has resulted in the production of tremendous amounts of transcript data associated with each gene. In turn, this data has revealed the existence of many different transcript isoforms associated with each gene. Given that control of gene expression via transcriptional regulation is key to our understanding of both developmental and disease states, we must ask if these different isoforms are the signal or the noise when evaluating the regulation of protein-encoding transcripts. In other words, do transcript isoforms have a fundamental role in regulating the presence or absence of a given gene product? Do they always contain important regulatory signals? If the answer is, as we believe, “yes”, then being able to determine their individual contributions to gene expression/regulation is fundamental. Yet, because the majority of transcript isoforms share substantial regions of identity with their own respective canonical transcripts, the determination of their individual contributions to gene expression is difficult.

In this work, we have detected contradictory cases where quasi-identical transcripts were declared as both being repressed (Figs. 13 and 14) or activated (Figs. 15 and 16). We have also observed a high number of transcripts that were considered significant in an experiment-specific manner (see Figs. 6 and 7). Together, while we think that the set of genes we identified to be significant are indeed significant, we believe that our results underscore the urgent need for the development of algorithms that will evaluate the contribution of each individual transcript isoform to gene expression. There needs to be recognition that not all genes are equal, and that genes coding for a single transcript may need to be evaluated differently than those having multiple transcripts. In addition, it is important to recognize and address the fact that many transcripts have perfect identity with transcripts produced by different loci. We believe that the critical examination of existing algorithms, along with the development of new ones, will allow us to dissect and understand the complexity of gene regulation that is so critical for our understanding of development and disease.

## Acknowledgments

This work was in part performed using the Supercomputers Grace and Terra, managed and maintained by the Texas A&M University High Performance Research Computing Resources Center (http://hprc.tamu.edu). In particular, we would like to acknowledge Dr. Michael Dickens for his invaluable help and support during the realization of this work and Tatiana Aramayo for her invaluable help editing the manuscript.

## SUPPLEMENTAL DATA

The figures, tables, supplemental figure, supplemental tables, mapping commands, and analysis com- mands, corresponding to this work have all been deposited in the Zenodo public database, with a DOI 10.5281/zenodo.7055085.

